# A cryptochrome photoreceptor controls animal light-dependent growth and lifespan via evolutionary conserved hormonal pathways

**DOI:** 10.1101/2025.03.12.642703

**Authors:** Gabriele Andreatta, Federico Scaramuzza, Aida Ćorić, Lukas Orel, Kristin Tessmar-Raible

## Abstract

Natural light is severely affected by human impact on Earth, yet little is known about the roles light receptors have outside vision and rhythmic processes. Here we show that loss-of-function of the *light-receptive cryptochrome* (*l-cry*) in marine bristleworms significantly increases lifespan and adult size, similarly to wild-types reared in constant darkness. Quantitative transcriptomics revealed hormonal players crucial for invertebrate and vertebrate sexual development and reproduction affected in *l-cry* mutants. These include *nr0b1/2*, ortholog of *dax-1* (*nr0b1*) and *shp* (*nr0b2*), long considered vertebrate novelties. Depending on moon-phase, *nr0b1/2* is up- or down-regulated in *l-cry* mutants. Matching the complex regulation, loss of *nr0b1/2* function partially recapitulates *l-cry* phenotypes. Molecularly, *Platynereis* Nr0b1/2 affects steroidogenic and other endocrine pathways, nuclear receptor signaling, and transcription factor orthologs, involved in sexual developmental, reproductive, and timing processes in other organisms. Thus, our study reveals profound effects of light on adult animal life-time, likely at least in part by conserved endocrine pathways involved in sexual maturation and reproduction in annelids and vertebrates.

## Introduction

The duration of life is influenced by many factors. The program that determines this, and more generally life-history traits (i.e. lifespan, adult size, growth rate, reproductive timing, fecundity), can be plastically reshaped by environmental changes (*1–3*). While food availability/intake, temperature, social interactions or circadian synchronization have been shown to influence lifespan and growth (*4–12*), the role of light itself does not readily come to mind as lifespan determinant in animals. Understanding the non-visual effects of light is, however, particularly relevant considering that changes to natural illumination are among the largest anthropogenic effects with already documented influences on human health and the environment (*13, 14*). The latter is increasingly recognized for marine ecosystems, whose integrity and prosperity are threatened by anthropogenic light changes (*13, 15, 16*).

Like many marine animals from corals to vertebrates, the physiology and behavior of the bristleworm *Platynereis dumerilii* is highly dependent on the timing, intensity and spectrum of the light that reaches its habitat, corresponding in nature to sun- and moonlight (*14, 17, 18*). Consistently, it possesses a repertoire of at least ten light-sensitive opsins and two light-sensitive cryptochromes (*14, 19–22*) (Genbank ID: GCA_026936325.1). The light receptor L-Cryptochrome (L-Cry) functions as a light interpreter, discriminating between sun- and moonlight, and providing key information on moon phase (*22–24*). In line with this, *Platynereis l-cry-/-* knockouts show altered reproductive rhythms and entrainment dynamics (*22, 23*).

In addition to this chronobiological phenotype, we observed that *l-cry-/-* knockout worms exhibit an increase in lifespan of more than 40% due to an extension of their immature (“youth”) and premature (“puberty-like”) stages, during which they exhibit most characteristics of an adult worm, except reproductive traits. In reference to classical literature, both immature and premature are atoke stages, while the onset of reproductive traits marks the epitoke stage. While these worms also attain significantly larger sizes, no morphological abnormality was found. Our molecular analyses revealed an altered endocrine system, with the single worm ortholog of vertebrate atypical nuclear receptors Dax-1 (Nr0b1) and Shp (Nr0b2) as a major target of L-Cry signaling. By disabling this nuclear receptor (lost from arthropod and nematode lineages), we unveiled molecular signatures of conserved functions between *nr0b1/2* and its vertebrate orthologs. Overall, our study reveals light-dependent effects of photoreceptor function on animal lifespan and growth, and provides novel insights into the hormonal pathways downstream of animal photoreceptors that contribute to sexual development and reproduction.

## Results

### *l-cry-/-* knockouts extend their lifespan and adult size

Phenotypic analyses performed in density-controlled conditions revealed that *l-cry-/-* knockouts prolonged the time required to achieve sexual maturity/reproduction from fertilization, compared to wild-type (wt) siblings (**Fig. 1A, B**; **fig. S1A**; **table S9**). Specifically, male and female mutants required on average 2.7 and 2.6 months longer to complete their life-cycle, respectively, with an average lifespan increase of ∼45% and ∼40% (**Fig. 1B**; **fig. S1A**). The remarkable lifespan extension was accompanied by a significant increase in final size (+9% and +7% in male and female average segments number, +90% and +50% in average body wet weight) (**Fig. 1C-E**; **fig. S1B, C**). The prolonged lifespan of *l-cry-/-* knockouts correlated with an initial delay in body growth (**Fig. 1F**; **fig. S1D-G**), and particularly germline development, compared to age-matched *l-cry+/+* worms (**Fig. 1G, H**; **fig. S1H-J**). The extended growth period of the of *l-cry-/-* (**Fig. 1B**) finally resulted in the larger size and body weight, as the worms continued to grow when wild-types had long entered reproduction.

**Figure 1:**
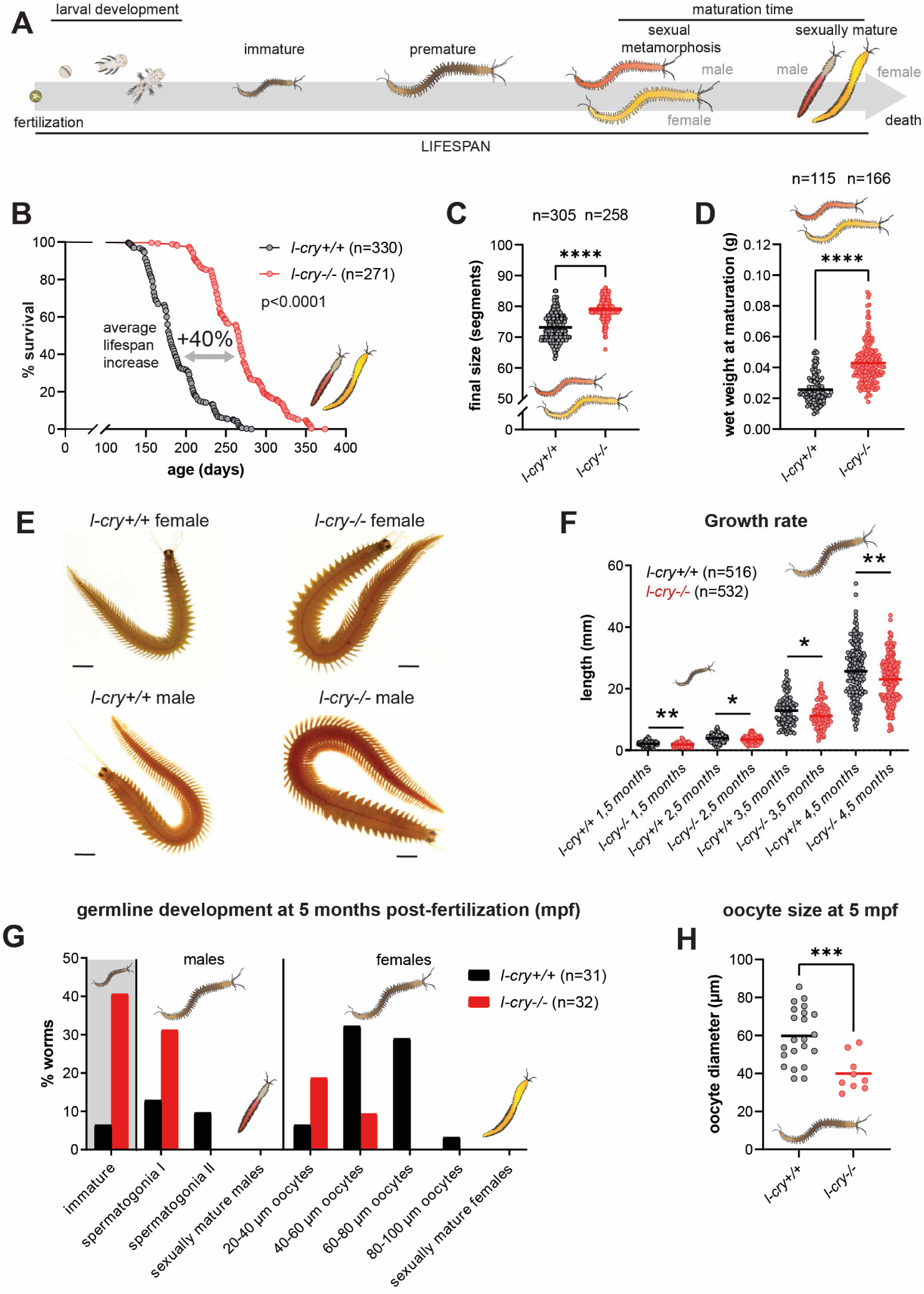
*l-cry-/-* knockouts show lifespan extension and increased final size. **(A)** Schematic representation of *P. dumerilii*’s development. For additional information: Supplementary text. **(B)** Survival (maturation) curves of *l-cry+/+* (black) and *l-cry-/-* (red) worms indicating the proportion of individuals that have yet to attain sexual maturity, and thus are still alive (% survival), over time. Lifespan: time-span between fertilization and reproduction/death. N ≥ 271 for each genotype. Log-rank (Mantel-Cox) test, p < 0.0001 (see **table S9**). **(C)** Final size of *l-cry+/+* (black) and *l-cry-/-* (red) worms as total number of segments at maturation. N ≥ 258 for each genotype. T test, **** p < 0.0001. **(D)** Wet weight of *l-cry+/+* (black) and *l-cry-/-* (red) worms during sexual metamorphosis. N ≥ 155 for each genotype. T test, **** p < 0.0001. **(E)** Representative pictures of *l-cry+/+* and *l-cry-/-* males and females. Scale bars at equal length in arbitrary units. **(F)** Growth rate of age-matched *l-cry+/+* and *l-cry-/-* worms measuring total worm length across immature and premature development. N ≥ 516 for each genotype. T test, * p < 0.05, ** p < 0.01. **(G)** Characterization of germline development in age-matched *l-cry+/+* and *l-cry-/-* worms after 5 months from fertilization. Bars represent the proportion found at each stage of germline development. N ≥ 31 for each genotype. **(H)** Maximum oocytes size (diameter) at 5 months from fertilization in *l-cry+/+* and *l-cry-/-* premature females. N ≥ 9 for each genotype. T test, *** p < 0.001.

We assessed two timepoints after the onset of reproductive age of wt siblings. While after 5 months post fertilization (mpf) ∼6% of the wt worms showed undifferentiated germline (immature worms: germline has not yet visibly differentiated into spermatogonia or eggs), this was the case in 40% of *l-cry-/-* worms (**Fig. 1G**). For animals that possessed differentiated spermatogonia or oocytes (premature worms, **Fig. 1A**), *l-cry-/-* worms showed less advanced stages of germline development compared to *l-cry+/+* individuals, quantified by measuring oocytes size in the two genotypes (**Fig. 1G, H)**. After 6 months post fertilization, ∼10% of knockouts were still immature compared to 0% of wt (**fig. S1H)**. Similar to the previous timepoint, *l-cry+/+* worms showed more advanced gametogenesis, with ∼60% that had already reached sexual maturity and had reproduced, compared to 0% in knockouts (**fig. S1H, I**). Despite significant differences in their growth rate, oocytes from both sexually mature *l-cry+/+* and *l-cry-/-* females reached comparable sizes at release (∼160-190 μm, **fig. S1J**). Given that knockout worms grow significantly beyond the age and size of wt worms without initiating sexual maturation, a delayed germline development is a plausible explanation.

### Wild-type animals exhibit lifespan extension and increased final size similar to *l-cry-/-* **worms under constant darkness**

As the phenotypes observed resulted from the lack of a functional photoreceptor, we next tested how lifespan and final size of *l-cry+/+* and *l-cry-/-* worms were affected by the exposure to naturalistic light conditions and altered light regimes (**Fig. 2A**).

**Figure 2:**
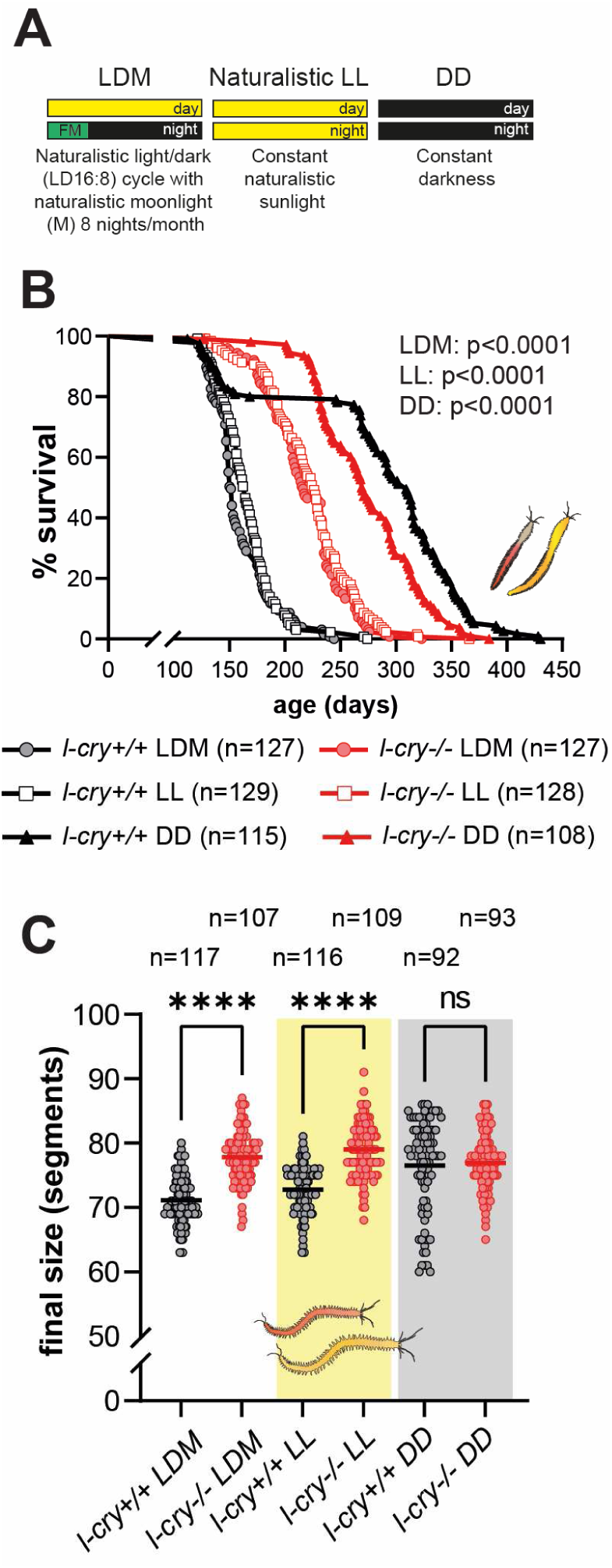
Lifespan and growth extension are L-Cry- and light-dependent. **(A)** Schematic representation of light conditions used. **(B)** Survival (maturation) curves of *l-cry+/+* (black) and *l-cry-/-* (red) worms over time. Lifespan was considered the time-span between fertilization and reproduction/death. Filled circles = light/dark (LD 16:8) + 8 nights of naturalistic moonlight; white squares = light/light (LL); filled triangles = dark/dark (DD). N ≥ 108 for each genotype/condition combination. Log-rank (Mantel-Cox) test, p < 0.0001 (see **table S9**). p values shown in B) refer to wt vs. *l-cry-/-* comparison under the respective mentioned condition. **(C)** Final size of *l-cry+/+* and *l-cry-/-* worms quantified as total number of segments at maturation. White background = light/dark (LD 16:8) + 8 nights of naturalistic moonlight; yellow background = light/light (LL); grey background = dark/dark (DD). N ≥ 92 for each genotype/condition combination. One-way ANOVA, **** p < 0.0001.

Animals were reared under standard laboratory settings and comparable density conditions for ∼2 months and subsequently transferred to one of the following conditions: i) a light/dark cycle using naturalistic sunlight and moonlight (LDM, (*22, 25*), ii) constant sunlight (LL), iii) constant darkness (DD, **Fig. 2A**). We reasoned that, if the lifespan/growth extension is the result of a delayed germline development, light-dependent effects on worm lifespan and growth should still be visible after the transfer at ∼2 months. Under both naturalistic LDM and LL conditions, *l-cry-/-* knockouts extended their lifespan and attained larger final size compared to wt siblings similar to the previous experiments conducted under standard artificial LDM conditions (compare **Fig. 1B, C**; **fig. S1A, B** vs. **Fig. 2B, C**; **fig. S2A, B**; **table S9**). In contrast, under DD, *l-cry+/+* animals significantly increased their lifespan and final size, reaching values comparable to those of *l-cry-/-* knockouts under identical conditions (**Fig. 2B, C**; **fig. S2A, B**; **table S9**). More specifically, we observed that under DD both *l-cry+/+* and *l-cry-/-* worms increased their lifespan compared to their respective counterparts reared under LDM and LL conditions (**Fig. 2B**; **fig. S2A**). Importantly, wt and mutant worms showed no significant difference in their final size under DD. For lifespan, the average difference is markedly reduced (3% compared to 34% and 37% for LL and LDM conditions, respectively) and caused by an even further increased lifespan of the wild-type. The latter is likely due to a complex interplay with the additional light receptors impacting on the worms’ physiology and changes in extrinsic factors, such as the microenvironment in the boxes (see Supplementary text for additional details and (*26*)).

Taken together, this evidence suggests that the lifespan and overall body growth extension observed under illuminated conditions are indeed majorly due to L-Cry’s light receptive function.

### The endocrine system, particularly *nr0b1/2*, the worm ortholog of vertebrate nuclear receptors *dax-1* and *shp*, is a target of L-Cry

To investigate the molecular mechanisms underlying the lifespan and growth alterations of *l-cry-/-* knockouts, we bulk-sequenced head transcriptomes of *l-cry-/-* mutant and wt worms. The worm brain has been shown to orchestrate maturation/reproductive timing (*22, 23, 27–30*), and to modulate both germline development and posterior segment addition (i.e. body growth) (*29, 31–34*). L-Cry localizes prominently into the brain, specifically to the eyes and a region that exhibits functional and molecular similarities to the vertebrate hypothalamus (*22, 27, 35, 36*).

We sequenced head transcriptomes from immature and premature female worms (**Fig. 3A**). We focused on female worms, because their gonadal stages can be better differentiated than male worms. In immature worms at ∼30-45 segments, germline proliferation and early differentiation occur (*34, 37*), while in premature females, more advanced processes of germline development and gametogenesis can be quantitatively assessed. In males, such temporal germline analyses are less precise as spermatogonia do not grow progressively in size till sexual maturity (*38*).

**Figure 3:**
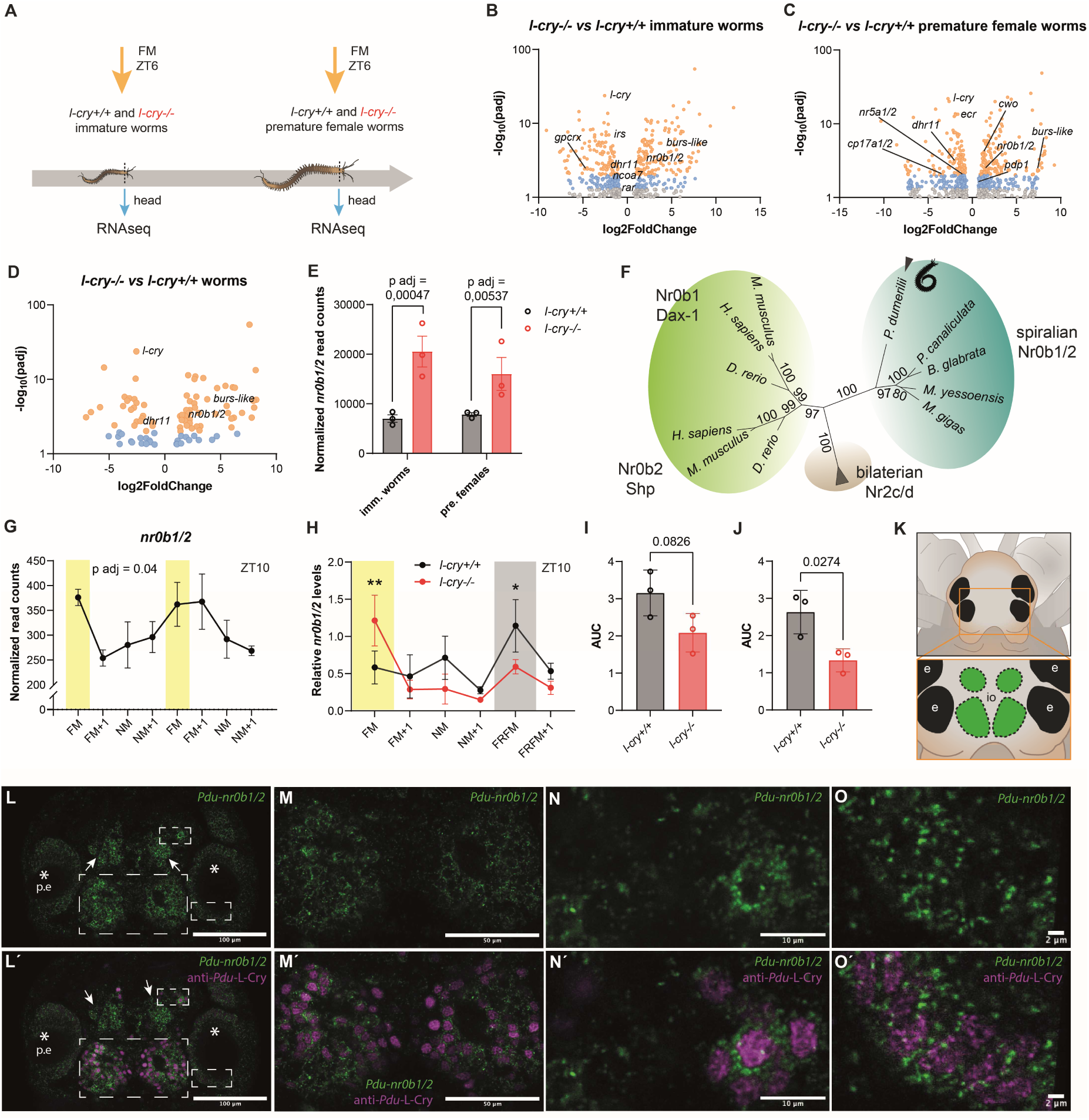
The endocrine system, particularly the *nr0b1/2-Platynereis* ortholog of vertebrate *dax-1/shp* nuclear receptors-, is a target of L-Cry. **(A)** Schematic representation of the sampling scheme used in (B) and (C). FM = full moon, ZT = Zeitgeber Time. **(B, C)** Volcano plots summarizing major expression changes between *l-cry+/+* and *l-cry-/-* immature (B) and premature females (C) worms of transcripts encoding (putative) components of the endocrine system (+ *l-cry*), one of the most regulated processes in *l-cry-/-* knockouts. To display the relationship between adjusted p value (padj) and Fold Change, these values have been converted to -log_10_(padj) and log2FoldChange, respectively. For the sake of clarity only transcripts with padj ≤ 0.1 are displayed. Orange dots: padj ≤ 0.01; blue dots: padj ≤ 0.05. **(D)** Volcano plot visualizing only the transcripts commonly regulated in both immature and premature female worms, with a particular emphasis on the endocrine system. Orange dots: padj ≤ 0.01; blue dots: padj ≤ 0.05. **(E)** Normalized *nr0b1/2* read counts from *l-cry+/+* (black) and *l-cry-/-* (red) immature and premature female worms. T test. **(F)** Phylogenetic tree reconstruction of bilaterian Nr0b1/2 nuclear receptors. Nr2c/d nuclear receptors were used as outgroup. **(G)** Normalized *nr0b1/2* read counts at ZT10 in *l-cry+/+* premature worms over the course of two lunar months. FM = full moon; FM+1 = FM+1 week; NM = new moon; NM+1 = NM+1 week. Yellow rectangles: FM phases. RAIN adjusted p value (p adj) = 0.04. **(H)** Relative *nr0b1/2* levels at ZT10 in *l-cry+/+* (black) and *l-cry-/-* (red) immature worms in the different lunar phases and in free-running (FR) conditions. FM = full moon; FM+1 = FM+1 week; NM = new moon; NM+1 = NM+1 week; FRFM = free-running full moon; FRFM +1 = FRFM+1 week. The yellow rectangle: FM phase, grey rectangle: subjective FM period (FRFM), where the moonlight stimulus was not provided. Two-way ANOVA. * p < 0.05; ** p < 0.01. **(I)** Quantification of overall *nr0b1/2* expression levels as Area Under the Curve (AUC) in *l-cry+/+* and *l-cry-/-* immature worms in all lunar phases shown in (H). T test. **(J)** Quantification of overall *nr0b1/2* expression levels as Area Under the Curve (AUC) in *l-cry+/+* and *l-cry-/-* immature worms in all lunar phases shown in (H), excluding the FM phase, the only timepoint in which *nr0b1/2* was upregulated in *l-cry-/-* compared to *l-cry+/+* worms. T test. **(K)** Schematic representation of the worm’s head, with a focus on the brain regions primarily expressing L-Cry and *nr0b1/2* (in green). e = eye; io = infracerebral organ. **(L-O)** Hybridization Chain Reaction (HCR)-based mRNA staining showing brain areas expressing *nr0b1/2*. (M-O) show higher magnifications of dotted rectangles drawn in (L). **(Ĺ-Ó)** Combined HCR showing brain areas expressing *nr0b1/2* (green) with L-Cry protein immunolabelling (magenta). (M’-O’) show higher magnifications of dotted rectangles drawn in (L’). * = eye; p.e. = posterior eye. Scale bars are 100, 50, 10, 2 μm, respectively.

*l-cry-/-* knockouts showed altered expression levels of transcripts associated with a variety of processes (**table S1**). However, what stood out was a broad regulation of the worm’s endocrine system in both immature and premature stages (**Fig. 3B, C**). Specifically, the lack of L-Cry signaling resulted in the altered expression of a *bursicon* (*burs*)-like glycoprotein hormone, *Nuclear Receptor subfamily 0 group B 1/2* (*nr0b1/2*), and the steroidogenic enzyme *Dehydrogenase/reductase SDR family member 11* (*dhr11*) in both developmental stages (**Fig. 3B-D**; see **fig. S15**; **Fig. 3E, F**; **fig. S6**, respectively). In addition to these generally differentially regulated transcripts, we identified a variety of hormonal players regulated in a stage-dependent manner, including insulin receptor substrates (*irs, als*), putative G protein-coupled receptors (*GPCRs*), nuclear receptors like *Retinoic acid receptor* (*rar*), *Ecdysone Receptor* (*ecr*), *Nuclear Receptor subfamily 5 group A1/2* (*nr5a1/2*), *Vitamin D3 receptor-like* (*vdr-like*), and *Ecdysone-induced protein 78C-like* (*e78c-like*) (**fig. S12**; **fig. S17**), components of thyroid hormone signaling (i.e. diodinase enzyme, *dio*), and putative steroidogenic enzymes like *Steroid 17-alpha-hydroxylase/17,20 lyase-like* (*cp17a1/2*) and *cholesterol 24-hydroxylase* (*cp46a1*) (see **fig. S7, 8**) (**Fig. 3B, C**; **table S1**). Orthologs of these factors coordinate developmental and reproductive aspects in either arthropods (i.e. *als, irs, burs* family, *ecr, e78c, hr96*) (*39–52*) or vertebrates (*nr0b1, nr0b2, dhr11, nr5a1, dio, cp17a1, rar, vdr*) (*53–66*). In mammals, CP17A1 and CP46A1 are key enzymes for corticoid/sex steroid and cholesterol/oxysterol biosynthesis, respectively (*66, 67*). Moreover, we found among the most regulated transcripts, *PAR domain protein 1* (*pdp1*) and *clockwork orange* (*cwo*) (**Fig. 3C**; **table S1**) two transcriptional regulators mostly known for their function in the circadian clock of *Drosophila melanogaster*, and at first glance mainly consistent with *l-cry’s* role in circadian-circalunidian timing (*68–71*). Yet, they also represent homologs of vertebratés PAR subfamily of basic leucin zipper (PAR bZip) transcription factors *hepatic leukemia factor* (*hlf*), *thyrotroph embryonic factor* (*tef*), *albumin D-box binding protein* (*dbp*), and *Differentiated Embryo-Chondrocyte expressed gene 1-2* (*dec1-2*) (**fig. S9**; **fig. S10**). Of note, mice deficient for HLF, TEF, and DBP display a dramatically shortened lifespan (*72*), whereas their common *Platynereis* ortholog, *pdp1*, is significantly upregulated in the long-lived *l-cry-/-* knockouts (**Fig. 3C**; **table S1**). Moreover, in insects, *pdp1* plays important roles in developmental progression, growth and reproductive arrest (*73–76*).

One of the most robustly regulated transcripts in *l-cry-/-* knockouts is the worm ortholog of vertebrate *Nuclear Receptor subfamily 0 group B* (*nr0b*) members (**Fig. 3B-F**): *nr0b1* (also *dax-1*) and *nr0b2* (also *shp*).

DAX-1 is an atypical orphan nuclear receptor crucial for the development of the hypothalamic-pituitary-gonadal (HPG) axis (*77*), steroidogenic tissues (gonads and adrenal glands) (*53*), gonads differentiation and maturation (*53*). SHP, besides roles in the regulation of bile acids, cholesterol and glucose metabolism in the liver (*78*), is also implicated in gonadal development and steroidogenesis (*54, 79*). *nr0b1/2* genes have long been considered a vertebrate novelty (*80*), primarily because they are absent from the ecdysozoan lineage, including fruit flies and nematodes (*81–83*) (**Fig. 3F**; **fig. S21**). Instead, our phylogenetic analysis revealed that *dax-1* and *shp* are likely the result of a vertebrate-specific gene duplication of a single ancestral *nr0b1/2* gene (**Fig. 3F**; **fig. S21**). Gene expression analyses by qPCR confirmed the upregulation of *Pdu-nr0b1/2* during full moon (FM) in *l-cry-/-* vs*. l-cry+/+* (**Fig. 3H-J**), while it was downregulated or less impacted at other lunar phases (**Fig. 3H**). Transcriptome sequencing of wt heads across different lunar phases confirmed a significant lunar regulation (**Fig. 3G**). Given the prominent role of *Dax-1* in the temporal control of ovulation in mammals, which ultimately dictates reproductive cycle length (*84*), it is noteworthy that L-Cry and the lunar cycle impact on the levels of its ortholog. Is it possible that pathways of reproductive timing share conserved molecular components downstream of environmental signal sensors?

Consistent with this hypothesis, we find that *nr0b1/2* mRNA prominently co-localized to cells demarcated by L-Cry antibody in the median neurosecretory brain regions and in the eyes (**Fig. 3K-O’**). All L-Cry^+^ cells possessed *nr0b1/2* mRNA (see **Fig. 3L-O’**), while a median-anterior region only possessed *nr0b1/2* mRNA (see arrows in **Fig. 3L, L’**). As we did not find any L-Cry^+^ cells without *nr0b1/2,* it suggests a likely tight functional interaction between the two factors, and that L-Cry can directly regulate *nr0b1/2* functions in the eye, posterior oval-shaped domain and a small medial anterior domain (**Fig. 3K-O’**).

A recent study in the clam *Tridacna crocea* indicates an involvement of *nr0b1/2* in spermatogenesis (*85*), and this, together with the findings presented above, prompted us to investigate the possible functions *nr0b1/2* might possess in *Platynereis* and how this might interconnect with the *l-cry* phenotypes. Given the complex temporal regulation of *Pdu-nr0b1/2* over the lunar cycle (**Fig. 3G, H**), and that depending on the phase of the lunar cycle it can be either up- or down-regulated in *l-cry-/-* worms **(Fig. 3H)**, it is obvious that none of the still relatively simple functional genetic manipulation available in *P. dumerilii* will fully mimic or produce opposite phenotypes of *l-cry* knockouts.

### *Platynereis nr0b1/2* mutants partially recapitulate *l-cry-/-* phenotypes

The *Platynereis nr0b1/2* transcript encodes a nuclear receptor with a protein length comparable to mammalian DAX-1 (rather than SHP) and other invertebrate Nr0bs (alignment **fig. S11**). By comparing vertebrate and spiralian sequences, the C-terminal putative ligand binding domain (LBD) shows overall higher similarity than the putative DNA-binding domain (DBD) (**fig. S11**). The typical LXXLL (or its variant LXXXL, (*86*), called here LXXLL-related) motifs, mediating DAX-1/SHP repressive activity on other nuclear receptors (*53, 87–89*), are present in putative DBDs and LBDs of both vertebrate and spiralian Nr0b1/2 proteins (**fig. S11**).

We generated *Pdu-nr0b1/2* mutants using established TALEN-targeted mutagenesis (*90*). We identified worms carrying 5-bp and 4-bp deletions, resulting in premature stop codons (**Fig. 4A**). As we did not identify *nr0b1/2* homozygous mutant worms reaching premature or mature stages under standard culturing conditions, we conducted a large-scale genotyping analysis of the progeny from *nr0b1/2+/-* incrosses at different stages of early development. We found Mendelian ratios of homozygous *nr0b1/2^Δ5/Δ5^* (hereafter referred as *nr0b1/2-/-*, likewise *nr0b1/2^Δ4/Δ5^* trans-heterozygous) worms at 1- and 6-days post fertilization (dpf) (**Fig. 4B**), suggesting fertilization and larval development were not impacted.

**Figure 4:**
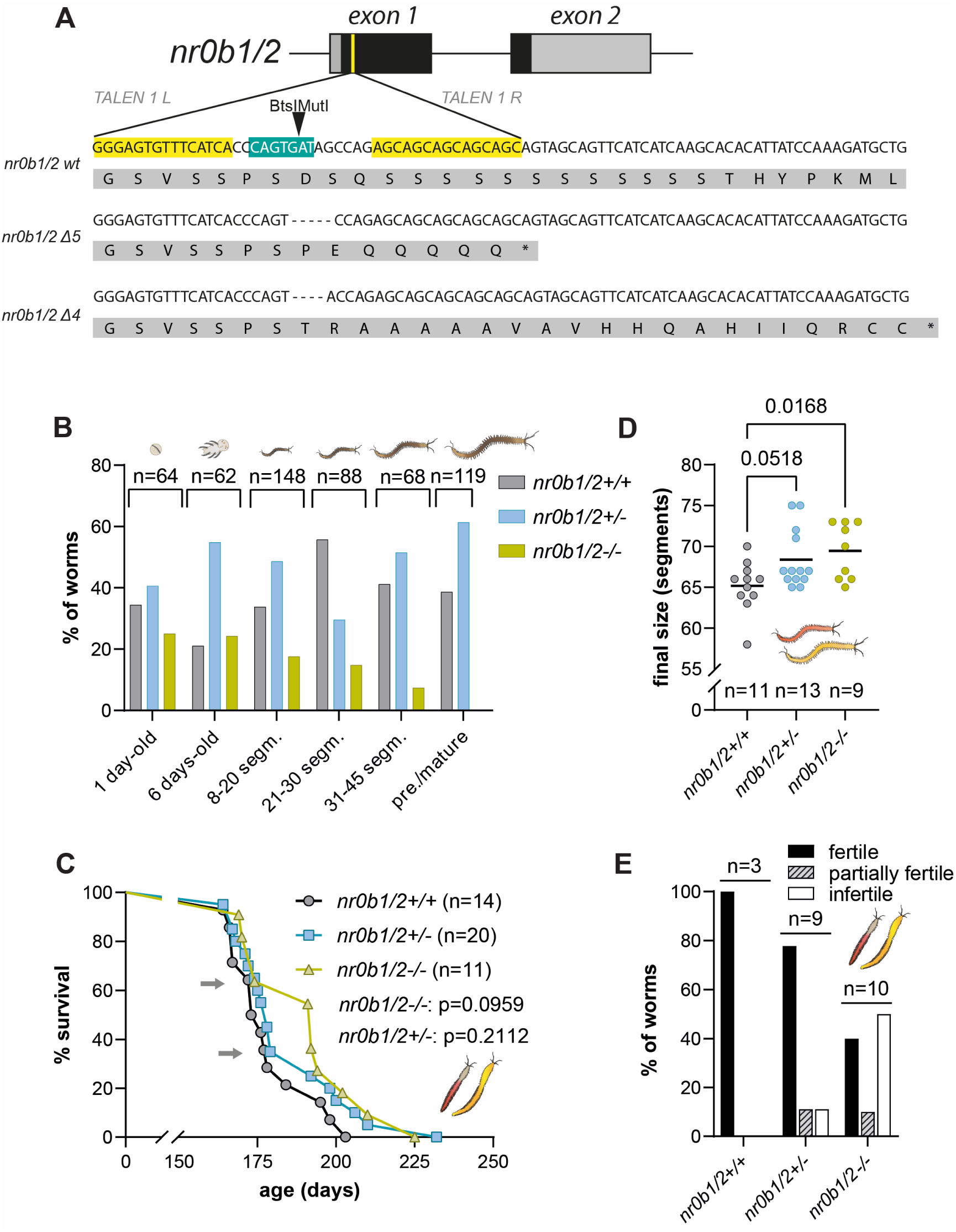
*nr0b1/2* knockout worms show fitness disadvantage and partially recapitulate the lifespan and growth extension of *l-cry-/-* mutants. **(A)** Schematic representation of *nr0b* locus highlighting the site in exon 1 selected for targeted mutagenesis and the alignment of both nucleotide as well as aminoacidic sequences of wild-type and the two frame-shift mutations identified (*nr0b1/2^Δ5^*and *nr0b1/2^Δ4^*). Rectangles represent exons (black: coding, grey: non-coding), yellow sections highlight the binding sites for TALENs, blue: restriction site around which TALENs were designed, white background: nucleotide sequences; grey: amino acidic sequences. (B) *nr0b1/2-/-* worms don’t survive under competitive conditions. Although present with frequencies approximating Mendelian distributions early in development, *nr0b1/2-/-* knockouts (light green) progressively reduce their proportion when reared in boxes together with *nr0b1/2+/-* and *nr0b1/2+/+* individuals, till their complete disappearance around the immature-premature boundary, when worms start differentiating their germline. Grey bars = *nr0b1/2+/+* animals; light blue bars = *nr0b1/2+/-* animals; light green bars = *nr0b1/2-/-* animals. N ≥ 62 for every timepoint. Chi-square test, df = 10, p < 0.0001. **(C)** Survival (maturation) curves of *nr0b1/2+/+* (black), *nr0b1/2*+/-(light blue) and *nr0b1/2*-/- (light green) worms indicating the proportion of individuals that have yet to attain sexual maturity, and thus are still alive (% survival), over time. N ≥ 11 for each genotype. Log-rank (Mantel-Cox) test to wildtype: p = 0.0959 (*nr0b1/2-/-*) and p = 0.2112 (*nr0b1/2+/-*) (see **table S9**). **(D)** Final size of *nr0b1/2+/+, nr0b1/2*+/- and *nr0b1/2*-/- worms quantified as total number of segments at maturation. N ≥ 9 for each genotype. One-way ANOVA. **(E)** Fertility assessment of *nr0b1/2+/+, nr0b1/2*+/- and *nr0b1/2*-/- animals displayed as % of fertile, partially fertile, or infertile worms. N ≥ 3 for every genotype. Chi-square test, df = 4, p < 0.0001.

Yet, from scoring genotypes of worms with ∼10-20 segments and onwards, we observed a progressive decline in the proportion of trans-heterozygous and homozygous *nr0b1/2-/-* animals and their subsequent disappearance after worms reached ∼30-40 segments (**Fig. 4B**). Consistently, under regular culturing conditions we never identified a *nr0b1/2-/-* worm larger than 34 segments. Of note, this phase of worm development coincides with the temporal window in which germ cell clusters should start proliferating (*37*). This process marks the beginning of a developmental stage requiring a more complex regulation of energy allocation (*30*), as a second major physiological program adds to body growth. Thus, we wondered whether lowering the competition with other genotypes would allow some homozygous *nr0b1/2* mutants to survive beyond 35-40 segments size, and potentially untill maturation. For this, we isolated and genotyped randomly-selected worms during early phases of worm development (∼10-30 segments, see **fig. S3E**).

Of the 45 worms, which survived the procedure, all individually-isolated *nr0b1/2-/-* worms matured into visibly distinct males and females. We next assessed lifespan, final size at reproduction and monthly spawning distribution (the major phenotypes of *l-cry-/-* knockouts) in the few *nr0b1/2-/-* we successfully raised for the entire life-cycle. *nr0b1/2*-/- worms exhibited a significantly increased size and also a clear trend to a longer lifespan compared to respective *nr0b1/2+/+* controls (**Fig. 4C, D**; **fig. S3A-C**; **table S9**) and phenotypically reminiscent of *l-cry-/-* animals (**Fig. 1B, C**; **fig. S1A, B**).

*Nr0b1/2* mutants showed no obvious morphological abnormalities in soma, germline/gametes and egg area (**fig. S3D**) and also showed no change in the monthly spawning pattern (**fig. S3G-I**).

This suggests that despite lunar regulation of *nr0b* transcripts in wt animals, *nr0b1/2* is rather involved in growth and possibly aspects of gonadal development, but not essential for its temporal monthly control. Given DAX-1/SHP roles in gametogenesis (*53, 54, 79*), we tested fertilization success in *nr0b1/2* animals reaching sexual maturation in isolated conditions by crossing them with a wt individual of the opposite sex, and evaluating the proportion of living/swimming larvae generated in the following 1-2 days. We found a significant reduction in the reproductive success of *nr0b1/2-/-* animals (both males and females), as 50% of their mating resulted in a complete absence of living larvae and presence of dead eggs, a clear indication of fertilization failure (**Fig. 4E**), while fertilization success of *nr0b1/2+/-* worms was mildly affected (**Fig. 4E**). Given that *nr0b1/2*-/- zygotes and larvae develop successfully, at least initially, when derived from heterozygous incrosses (**Fig. 4B**), our observations point at developmental defects in the germline of *nr0b1/2-/-* animals that are not morphologically visible. The analyses of *nr0b1/2* knockouts provide evidence that, consistent with its expression pattern and complex regulation, L-Cry signaling is partially transduced (directly or indirectly) via changes in *nr0b1/2* and its downstream network.

### *nr0b1/2* regulates endocrine pathways and transcriptional regulators associated to sexual development/reproduction and circadian rhythms, partially overlapping with L-Cry targets

In order to test which parts of the L-Cry signaling might be molecularly transduced via (direct or indirect) changes in *nr0b1/2* expression, we performed RNAseq analyses on *nr0b1/2* mutants and compared them to the corresponding *l-cry* results. We adopted two parallel strategies to cope with the problem of raising *nr0b1/2-/-* worms. First, in order to guarantee the sampling of *nr0b1/2-/-* worms, we collected 315 immature worms (12-40 segments, see **fig. S3F**) from several *nr0b1/2+/-* crosses (both *nr0b1/2^+/Δ4^* and *nr0b1/2^+/Δ5^*) over an 8 h-window. For each worm, anterior portions (including the head) were collected for RNAseq (**Fig. 5A**), and 6 biological replicates (BRs) generated by pooling worms with the same genotype (**Fig. 5A**). Second, we also sequenced the head transcriptome of previously genotyped immature *nr0b1/2+/+* and *nr0b1/2+/-* sampled at ZT10, a strategy which provided a higher diel temporal precision (**Fig. 5D**).

**Figure 5:**
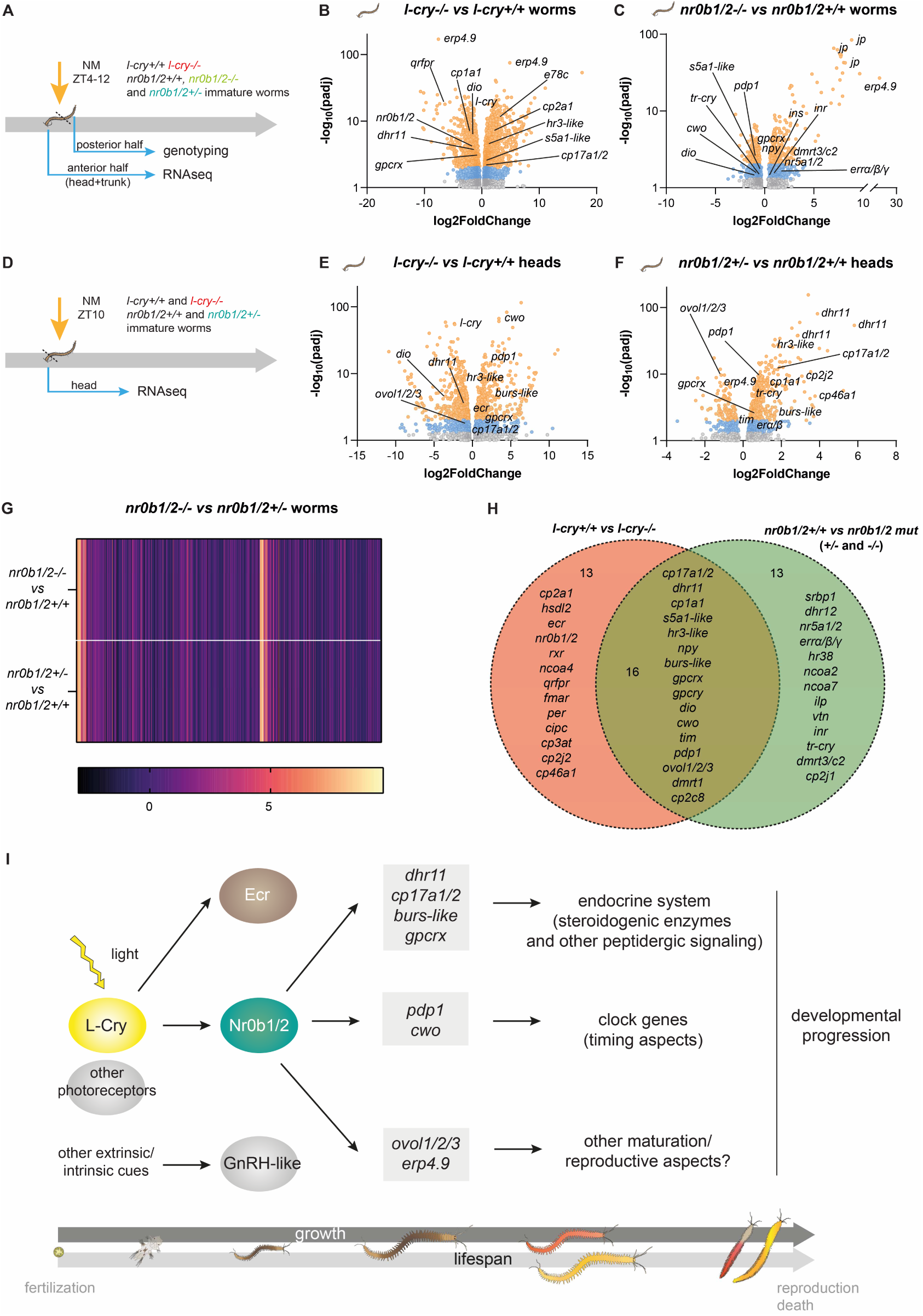
*nr0b1/2* regulates endocrine pathways and transcriptional regulators associated to sexual development/reproduction and chronobiology, partially overlapping with L-Cry targets. **(A, D)** Schematic representations of the sampling schemes used in (B, C), and (E, F), respectively. NM = new moon, ZT = Zeitgeber Time. **(B, C, E, F)** Volcano plots summarizing major expression changes between *l-cry+/+* and *l-cry-/-* immature worms (B), *nr0b1/2+/+* and *nr0b1/2-/-* immature worms (C), *l-cry+/+* and *l-cry-/-* immature worm heads (E), or *nr0b1/2+/+* and *nr0b1/2+/-* immature worm heads (F), with a particular emphasis on transcripts encoding (predicted) components of the endocrine system. To display the relationship between adjusted p value (padj) and Fold Change, these values have been converted to -log_10_(padj) and log2FoldChange, respectively. For the sake of clarity only transcripts with padj ≤ 0.1 are displayed. Orange dots: padj ≤ 0.01; blue dots: padj ≤ 0.05. **(G)** Heat map displaying similarities in molecular signatures arising from *nr0b1/2+/+* vs *nr0b1/2-/-* and *nr0b1/2+/+* vs *nr0b1/2+/-* comparisons. Every vertical line corresponds to the fold change of the same transcript in both analyses. **(H)** Venn diagram showing transcripts significantly and jointly regulated in both *l-cry+/+* vs *l-cry-/-* and *nr0b1/2* wt vs *nr0b1/2* mutants, focusing on those related to endocrine pathways, sexual development/reproduction, and biological rhythms.**(I)** Graphical summary of the L-Cry/Nr0b1/2 molecular network contributing to lifespan and growth in *Platynereis*. L-Cry mediates light effects by controlling *nr0b1/2* expression dynamics, as well as other important endocrine and non-endocrine pathways. Nr0b1/2, in turn, orchestrates the expression of downstream genes encoding key components of steroidogenic pathways (regulated also in *l-cry-/-* worms). Other light-dependent (i.e. other photoreceptors) or light-independent signaling pathways also impact on determining worm’s lifespan and growth dynamics.

To facilitate the identification of commonly regulated genes in *l-cry* and *nr0b1/2* mutants, we included *l-cry-/-* knockouts and wild-type counterparts sampled with the same strategies. We opted to perform our sampling during new moon (NM), as at this timepoint *nr0b1/2* expression in *l-cry+/+* and *l-cry-/-* worms shows a large difference (**Fig. 3H**). In both transcriptomic strategies, we confirmed the prominent regulation of the endocrine system in *l-cry-/-* knockouts (including genes involved in sexual development and reproduction in other animals) (see **table S2**; **table S3**). More specifically, we identified nuclear receptors in addition to *nr0b1/2* (*ecr* [**fig. S17**], *Nuclear hormone receptor HR3* [*hr3*]*-like* [**fig. S17**], *Retinoic acid X receptor* [*rxr*], *Nuclear receptor coactivators* [*ncoa2,4,7*]), peptidergic factors (*Pyroglutamylated RFamide peptide receptor* [*qrfpr*], *FMRFamide receptor* [*fmar*], *Pro-neuropeptide Y* [*npy*] (*21, 91*), a *bursicon* (*burs*)*-like* neuropeptide [**fig. S15**], other *GPCRs*), enzymes putatively involved in steroidogenesis (*cp17a1/2* [**fig. S7**], *dhr11* [**fig. S6**], *cp1a1, cp2a1, hydroxysteroid 11-beta-dehydrogenase 1 protein* [*dhi1l*], *hydroxysteroid dehydrogenase-like protein* [*hsdl2*], *3-oxo-5-alpha-steroid 4-dehydrogenase 1* [*s5a1*]*-like* [**fig. S14**]), and transcriptional regulators involved in rhythmic processes (*pdp1* [**fig. S9**], *cwo* [**fig. S10**], *period* [*per*], *CLOCK-interacting protein circadian* [*cipc*]) and sexual development (*transcriptional regulator ovo-like 1/2/3* [*ovol1/2/3*] [**fig. S18**], *doublesex- and mab-3-related transcription factor 1* [*dmrt1*] [**fig. S16**]), with several prominently regulated during FM as well (*dio, cp17a1/2, dhr11, ecr, cwo, pdp1, nr0b1/2*) (**Fig. 3B, C**; **Fig. 5B, E**; **table S1**; **table S2**; **table S3**). Of note, we also found *nr0b1/2* significantly downregulated at NM in *l-cry-/-* knockouts, consistent with the qPCR data (see **Fig. 3H**; **Fig. 5B**).

From the analyses of *nr0b1/2-/-* and *nr0b1/2+/-* transcriptomes, we noticed that (with one single exception (*ependymin-related protein 4.9*, [*erp4.9*]), there were no significant differences between the two genotypes at the molecular level (see **Fig. 5G**; **table S3**), consistent with their overall phenotypic similarity (**fig. S3A-I**). This likely suggests compensatory effects in the surviving homozygous animals. Differential expression analyses between each transcriptome and its *nr0b1/2+/+* counterpart showed a significant regulation of similar gene classes identified in *l-cry-/-* worms, including nuclear receptors and coactivators (*nr5a1/2* [**fig. S12**], *estrogen-related receptor α/β/γ* [*errα/β/γ*] [**fig. S13**], *hr38-like* [**fig. S17**], *ncoa2, ncoa7*), putative steroidogenic enzymes (*s5a1-like* [**fig. S14**]), cytochromes P450 (i.e. *cp2j1, cp2c8*), neuropeptides (*npy*, an *insulin-like peptide* [*ilp*], *prepro-vasotocin-neurophysin* [*vtn*]), peptide receptors (*insulin receptor* [*inr*], other *GPCRs*), thyroid hormone signaling (*dio*), transcriptional regulators involved in sexual development (*doublesex- and mab-3-related transcription factor 3/C2* [*dmrt3/c2*] [**fig. S16**]), rhythmic processes (*pdp1, cwo, vertebrate-like cryptochrome* [*tr-cry*]), and cholesterol biosynthesis/lipid homeostasis (*sterol regulatory element-binding protein 1* [*srbp1*]) (**Fig. 5C**; **table S3**). Several genes or pathways were represented in *nr0b1/2+/-* as well (*nr5a1/2, ncoa2, s5a1-like, npy, ilp, GPCRs, dio, dmrt1, dmrt3/c2, cp3ab, cp2cn*) (**fig. S4C**; **table S3**).

Using the second strategy (sampling at a precise timepoint), we complemented the transcriptional landscape affected by Nr0b1/2 signaling, including a prominent regulation of putative steroidogenic enzymes (*dhr11* [**fig. S6**], *cp17a1/2* [**fig. S7**], *cp1a1, dhr12* [**fig. S6**], *17-beta-hydroxysteroid dehydrogenase 13* [*dhb13*]), cholesterol homeostasis (*cp46a1* [**fig. S8**]), other cytochromes P450 (*cp2j2, cp3at*), nuclear receptors (*hr3-like* [**fig. S17**], *Vitamin D3 receptor-like* [*vdr-like*] [**fig. S17**], *Estrogen receptor* [*erα/β*]), peptidergic signaling (*bursicon* (*burs*)*-like* and *gpcrx*, a still unidentified GPCR [see **table S5**, see Supplementary text]), and again transcriptional regulators involved in both rhythmic processes (*pdp1* [**fig. S9**], *tr-cry, timeless* [*tim*]), and sexual development (*ovol1/2/3*, [**fig. S18**]) (**Fig. 5F**; **table S2**).

Among the most regulated transcripts between *nr0b1/2* mutants and *nr0b1/2+/+, dhr11* (**fig. S6**), *cp17a1/2* (**fig. S7**), *s5a1-like* (**fig. S14**), *nr5a1/2* (**fig. S12**), *errα/β/γ* (**fig. S13**), *dmrt1, dmrt3/c2* (**fig. S16**)*, ovol1/2/3* (**fig. S18**) are all homologs (and in some cases single worm orthologs) of important components of the vertebrate molecular machinery orchestrating steroidogenesis, sexual development, and/or directly interacting with DAX-1 (*53, 55, 59, 66, 88, 92–95*). In mammals, *DHR11, CP17A1* and *S5A1* are involved in the biosynthesis of steroid hormones, particularly of sex steroid hormones (*55, 66, 94*). Yet, it has been proposed that steroidogenesis evolved independently in the three major bilaterian linages (Vertebrata, Spiralia, Ecdysozoa) (*96*). In particular, no genes orthologous to vertebrate steroidogenic enzymes were thought being present in lophotrochozoans (part of the Spiralia clade that includes annelids) (*96*). While our analyses do not show that the conserved enzymes identified take part to the very same reactions in annelids, our findings challenge the notion of a completely independent evolution of steroidogenesis between deuterostomes and protostomes. It suggests rather secondary losses in specific lineages (i.e. ecdysozoans), as observed for several other genes (*83, 97, 98*).

We then intersected the *nr0b1/2* datasets with the respective *l-cry* ones to identify commonly regulated genes (**Fig. 5H**). By intersecting candidates differentially expressed in *l-cry-/-* and *nr0b1/2-/-* animals (relative to their wt counterparts, first strategy, see **Fig. 5A**), we found peptidergic GPCRs (*gpcrx, gpcry*, **table S5, table S6**, see Supplementary text), the nuclear receptor coactivator *ncoa2*, the putative steroidogenic enzyme *s5a1-like*, and the cytochrome P450 *cp2c8* (**fig. S4A, F**; see **table S3, 4**). On the other hand, intersecting candidates differentially expressed in *nr0b1/2+/-* and *l-cry-/-* heads (second strategy, **Fig. 5D**), we found putative steroidogenic enzymes (*dhr11, dhr12, cp17a1/2*), other cytochromes P450 (*cp3at, cp2j2*), the nuclear receptor *rxr*, a *bursicon* (*burs*)*-like* neuropeptide, *gprx*, and the transcriptional regulators *pdp1* and *ovol1/2/3* (**fig. S4B, G**; see **table S2, 4**). Gpcrx shares a robust similarity with Allatostatin A receptors (AstAR) and Gonadotropin-releasing hormone receptors (GnRHR), by reciprocal best blast against other invertebrate genomes (**table S5**, see Supplementary text). Although we could not pinpoint the exact phylogenetic relationships for this GPCR, both classes identified share important roles in development/maturation and reproduction in invertebrates and vertebrates (*30, 99*).

All combined, our data suggest that light-dependent phenotypes of *l-cry* knockouts (control of lifespan and growth but not monthly reproductive timing) are partially mediated by *nr0b1/2*, likely through the transcriptional regulation of a specific set of endocrine players and transcription factors (**Fig. 5H, I**) which orchestrate sexual development and maturation in both vertebrates and invertebrates.

## Discussion

The present study emphasizes the strong and direct impact of light on processes that are typically not immediately associated with light-dependence in animals: lifespan and growth.

As the regulation of lifespan and growth are relevant for species fitness, our findings are also ecologically relevant. Yet, what could be the reasons for such control in the wormś natural habitat? In the ocean, light intensity changes significantly in relation to depth and time (*14, 21, 100, 101*). Thus, the light-dependent control of lifespan and growth trajectories might evolve around a trade-off between reproductive success and predator avoidance (**fig. S5**). In many species (including *Platynereis*) larger bodies can accommodate a higher number of gametes (as the latter show comparable size between *l-cry+/+* and *l-cry-/-* worms), hence likely increasing reproductive success of a given individual. Consistently, the offspring numbers of *l-cry-/-* animals are typically at least twice as large as wild-type. Yet, larger animals take more time and resources to grow and are more easily noticed by predators, particularly when inhabiting shallow waters. In this context, a light-sensitive developmental modulation could enable worms to maximize both reproductive success and survival, adjusting their size (and thus number of gametes) based on lighting conditions (and thus risk of predation) (**fig. S5**). An increased number of offsprings also allows for wider progeny dispersal and by this can increase habitat diversity and reduce intraspecific competition (**fig. S5**). In line with this hypothesis and even though *P. dumerilii* lives primarily in 1 to 10 meters-deep shallow waters (*18*), individuals of this species have been reported and collected even up to 100 meters depth (*102, 103*).

Light information in the worms’ habitat also varies in a seasonal-dependent manner not only in terms of photoperiod, but also relative spectral intensities (*21*). The adjustment of lifespan and growth in a light-dependent manner could thus also be part of a seasonal optimization program reminiscent of those described for insects and vertebrates that enable animals to time their reproduction (or specific developmental transitions) to likely favorable environmental conditions (*104, 105*).

The phenomena we describe are unlikely to represent *Platynereis* peculiarities, as light-dependent modulation of developmental time has been shown in flies (*106*) (with longevity co-varying with this life-history trait, see (*107*)). However, unlike in flies, our study provides evidence for extensive molecular similarities in the reproductive machinery of annelids and vertebrates, often considered to have evolved largely independently. Thus, these light-dependent effects provide further insights into the possible implications of anthropogenic impacts on natural illumination, via light pollution, but also by reductions of light intensity caused by floating solar farms, as well as the increasingly discussed blocking of sunlight to prevent further global warming. Our work suggests that such interventions pose risks to environments that were so far unknown.

The prominent role of *Nr0b1* and *Nr0b2* (*Dax-1* and *Shp*) in the hormonal control of reproduction in vertebrates (*53, 54, 79*), recent functional evidence for the involvement of *nr0b1/2* in mollusc spermatogenesis (*85*), and its contribution to lifespan/growth control via conserved endocrine pathways, suggest that these are likely ancestral roles for this class of nuclear receptors. Of note, transcripts encoding putative steroidogenic enzymes are predominantly upregulated in *nr0b1/2* mutants, in line with the repressive role that DAX-1 exerts on the steroidogenic transcriptional landscape in mammals (*53*). Additionally, the nuclear receptors regulated in *nr0b1/2* mutants (i.e. *nr5a1/2, errα/β/γ*) suggest conserved regulatory signatures within *nr0bs* networks (*53*). *nr5a1/2* is the single worm ortholog of vertebrate *Steroidogenic Factor 1* (*sf-1, nr5a1*) and *Liver Receptor Homolog 1* (*lrh-1, nr5a2*), with the encoded proteins physically interacting with DAX-1 and SHP, which negatively modulate their nuclear receptor activity (*53, 108*). Specifically, DAX-1 is known to inhibit SF-1-mediated transcriptional activity, which targets genes involved in steroidogenesis, sexual development, and reproduction (*53*), including *Dax-1* itself (*109, 110*). In turn, SF-1 and LRH-1, promote *Shp* expression (*111*), creating multiple interconnected feedback loops.

An analogous regulatory scenario is in place also for mammalian ERRs, as this nuclear receptor induces *Dax-1/Shp* expression, which in turn inhibits the transactivation of the former (*112–114*). In *Platynereis, nr5a1/2* and *errα/β/γ* transcripts are all upregulated in *nr0b1/2* mutants, again potentially suggesting a repressive role of Nr0b1/2 on their expression. In line with Nr0b1/2 pivotal role in orchestrating aspects of sexual development also in *Platynereis*, among the regulated genes in *nr0b1/2* mutants we found homologs of *dmrt1* (and *dmrt3/c2*, also potentially involved in sexual development), and *ovol1/2/3*. Proteins sharing the DM domain (i.e. Doublesex and mab-3 related transcription factors 1, Dmrt1) are transcription factor known as conserved actors in male sex differentiation and gametogenesis in bilaterians (*92, 115, 116*), yet found to coordinate oogenesis as well (*117*). *Ovol1/2/3* encodes an evolutionary conserved zinc finger transcription factor which controls germline development in both flies (*118–120*) and mammals (*95, 121, 122*). In addition, we also identified signatures reminiscent of SHP functions not directly connected with sexual development/reproduction, like lipid/cholesterol metabolism (*cp46a1*), transcriptional control of other *CP450s* (i.e. *cp2j2, cp3at*), and the regulation of rhythmic processes (*pdp1, tr-cry, tim, rev-erbα/β/nr1d1/2*) (*123*). Indeed, in mammals, SHP orchestrates nutrient availability and lipid metabolism with the liver circadian clock, by regulating *BMAL1* (*124*), and interacting with RORγ and Rev-Erbα (*125*). Overall, this suggests that, after *dax-1* and *shp* emerged from an originally single gene as a result of vertebrate whole genome duplications, they underwent processes of sub-functionalization. Of course, we do not exclude that neo-functionalization processes also led to the evolution of novel roles in the two vertebrate genes, or that spiralian/species-specific functions were not inherited by either of the two vertebrate genes or evolved secondarily. An example of the latter might be the strong regulation of *jaw proteins* (*JPs*) in *nr0b1/2* mutant animals. However, our data largely support an as of yet unforeseen likely similarity between the downstream molecular pathways. Further future investigations will provide a better understanding of the details and implications of the evolutionarily conserved versus deviations in the different pathways.

## Acknowledgments

We are grateful to Andrij Belokurov, Margaryta Borysova and Netsanet Getachew for routine worm cultures and genotyping support, Birgit Poehn for providing worm head samples and RNA extracts, as well as all members of the Tessmar-Raible and Raible labs for constructive discussions. We thank the Next Generation Sequencing Facility at Vienna BioCenter Core Facilities (VBCF), member of the Vienna BioCenter (VBC), Austria for excellent services concerning all bulk sequencing.

## Funding

This work was supported by

Marie Curie Research Training Network: Project N° 101109842 (GA)

Helmholtz Society, distinguished professorship by the Alfred Wegener Institute Helmholtz Centre for Polar and Marine Research (KT-R)

H2020 European Research Council, ERC Grant Agreement #819952 (KT-R)

Austrian Science Funds (FWF), SFB F78 (KT-R)

Human Frontier Science Program (HFSP), #RGP021/2024 (KT-R)

None of the funding bodies was involved in the design of the study, the collection, analysis, and interpretation of data or in writing the manuscript.

## Author contributions

Conceptualization: GA, KT-R

Methodology: GA, FS, AC, LO, KT-R

Investigation: GA, FS, AC, KT-R

Visualization: GA, AC

Funding acquisition: KT-R

Project administration: KT-R

Supervision: GA, KT-R

Writing – original draft: GA, KT-R

Writing – review & editing: GA, FS, AC, LO, KT-R

## Data and materials availability

All data are available in the main text, and the supplementary materials (text and figures).

## Supplementary Materials

Materials and Methods

Supplementary Text

Figs. S1 to S22

Table S1 to S9

