## Supplementary Material for "A cryptochrome photoreceptor controls animal light-dependent growth and lifespan via evolutionary conserved hormonal pathways"

for

### Materials and Methods

Worm culture and phenotypic analysis

*Platynereis* cultures were maintained at 18°C (LD16:8) according to standard protocols as described previously (*30*). All animal work was conducted according to Austrian and European guidelines for animal research. For experiments assessing lifespan/maturation time, growth rate,final size, wet weight, and germline development (**Fig. 1G, H**; **fig. S1H-J**), ~1.5 months-old worms were collected and raised in boxes with the same density (0.075-0.085 worms/$cm^2$; ~0.04 worms/$cm^3$). Lifespan was defined the time span between fertilization and reproduction (spawning) and the subsequent death (which occurs shortly after gametes release). All details on statistical analyses of lifespan data are summarized in **table S9**. Final size was determined by counting the number of segments bearing parapodia in sexually mature (spawning) animals, since in this stage worms change their body morphology making worm length no more linearly predictive of their actual size. In all experiments assessing worm growth by segments count or length measurement, only worms with no signs of tail amputation/recent regeneration were considered. To measure the worms' wet weight, animals in the middle of sexual metamorphosis (bearing sexually dimorphic coloration but still exhibiting crawling behavior, and having no residual food in their gut) were anesthetized in a solution of 50% $MgCl_2$ and 50% sea water, and then gently dried on a lean-free tissue before being placed individually on glass slides and weighted. Worm´s wet weight was calculated as total weight minus the glass slide weight. Worm pictures were obtained using a Nikon SMZ18 stereomicroscope, DS-Ri2 camera and NIS Elements imaging software (Nikon) under bright field setting.

In the experiment designed to assess the effects of different light conditions on lifespan and final size, the selected conditions were: 1) naturalistic light/dark (LD) 16:8 (using naturalistic sunlight) with naturalistic moonlight (M) during 8 nights/month (LDM) (*22*, *25*); 2) constant sunlight (LL); 3) constant darkness (DD). To mimic naturalistic sunlight and moonlight conditions, we took advantage of NELIS (Natural Environmental Light Intensity System) (Marine Breeding Systems GmbH) (see (*22*, *25*)). Under LL, worms were not subjected to naturalistic moonlight, as this would have been covered by sunlight intensity during the subjective night. Light spectra were measured using an ILT950 Spectroradiometer (International Light Technologies) (see **fig. S20**), and oscillations in light intensity and temperature (°C) monitored using HOBO® devices (Onset Brands). The light intensity used was within the intensity range experienced by this species in its natural habitat (see (*21*)). For additional information on light spectra and intensity see (*25*). On average, temperature recorded in the three shelves was: LDM: 19.082 °C, LL: 19.081 °C, DD: 18.312 °C (see **table S8** for whole recordings). Worms were raised in standard LDM laboratory conditions (see (*22*, *126*)) for ~2 months since fertilization (~ for the entirety of their immature phase), and then transferred to one of the three conditions described above for the rest of their life-cycle (~ in proximity of their entry in premature phase, marked by germline proliferation and differentiation).

To assess growth rate, we first measured the length of 1.5 months-old worms kept in comparable density conditions and fed *ad libitum* with algae suspension. Then, worms were isolated in boxes with the same final density (see above) [**Fig. 1F**], or in 6 well-dishes [**fig. S1D, E**] (based to the experimental design), and fed normally with the same amount of food (spinach and Tetramin fish food). Every month, worms were anesthetized and photographed (**Fig. 1F**), or segments counted [**fig. S1D, E**]. Worm´s length was measured using Fiji v. 2.9.0 (ImageJ) (*127*), or assessed as number of segments (as clearly stated on graphs´ Y-axis). It has to be noted that, before the end of sexual metamorphosis, worm´s length and segment number co-vary linearly (see **fig. S19**). For regeneration experiments, worm´s trunks from age-matched ~30-40 segments-long immature individuals were halved, and the number of re-grown segments assessed every week by stereomicroscope inspection (after brief anesthesia) for 4 weeks. Animals were fed twice a week with a mixture of spirulina and Tetramin.

The status of germline development was evaluated for worms maintained in density-controlled conditions [**Fig. 1G, H**; **fig. S1H-J**] or 6 well- dishes [**fig. S3D**] (see above) using a Zeiss Observer Z1 microscope equipped with a CoolSNAP $HQ^2$ camera (Photometrics). Worms were anesthetized and two parapodia surgically removed from the middle part of their body. Germ cells were gently squeezed out and collected (see (*29*)). Pictures were taken at 20x magnification. The first timepoint selected for analyses of *l-cry* knockouts (after 5 months since fertilization) corresponded to the entry of the first worm into sexual metamorphosis, and we repeated the analyses after a month (6 months since fertilization), to provide a more dynamic temporal dissection of germline differentiation and maturation. Animals showing no discernable germline cells or still undifferentiated were considered immature worms. Oocyte diameter and area were calculated using the ZEN v. 3.1 software (Zeiss). For analyses of *nr0b1/2* mutants, we opted for an earlier assessment and a higher resolution because of the uncertainties about their survival till maturation.

For the qualitative assessment of fertility in *nr0b1/2+/+* (grey), *nr0b1/2+/-* (light blue), and *nr0b1/2-/-* (light green) worms (expressed as % of fertile/partially fertile/infertile mating), individuals (both males and females) from each genotype were mated with a wild-type counterpart to isolate male- or female-specific effects, and the offspring number visually assessed. Fertile = high offspring number (~> 50-100 larvae); partially fertile = very low offspring number (~10-20 larvae); infertile = no swimming/living larvae found. For females, the overall number of laid eggs (fertilized + unfertilized) was comparable.

Data were analyzed using Student's t-test or One-way ANOVA: * $p<0.05$; ** $p<0.01$; *** $p<0.001$; **** $p<0.0001$. Lifespan/age at reproduction was visualized and analyzed using survival curves and Log-rank (Mantel-Cox) test: * $p<0.05$; ** $p<0.01$; *** $p<0.001$; **** $p<0.0001$. Genotype frequency within batches and worm´s fertility were analyzed using $\chi^2$ test: * $p<0.05$; ** $p<0.01$; *** $p<0.001$; **** $p<0.0001$. Correlation between segment number and worm´s length was analyzed using simple linear regression. Statistical analysis was performed using Graph Pad Prism v. 9.

RNA sequencing

To investigate stage-specific transcriptional dynamics in *l-cry+/+* and *l-cry-/-* worms (see **Fig. 3A**), ~2 months-old immature worms were collected and decapitated at ZT6 during the full moon (FM) phase. Heads were snap-frozen in liquid nitrogen and stored at -80°C. For premature worms, individuals of comparable size from the two genotypes were collected three days before the sampling. Worms were anesthetized and two parapodia from the middle part of the body surgically removed. Germ cells were gently squeezed out and collected (*29*). Only females with clearly developed eggs were selected for further analyses, and maintained for two days in new boxes, prior decapitation and sampling at ZT6 (FM). All animals were starved ~7 days before sampling. For each biological replicate (BR) (3 BRs for each genotype/developmental stage combination), RNA was extracted from 8 immature worm heads or 4-5 premature female worm heads using the BioZym kit (Zymo Research). Similarly, in the experiment aiming at comparing transcriptional profiles in *nr0b1/2* (*nr0b1/2*+/+ and *nr0b1/2*+/-) and *l-cry* (*l-cry*+/+ and *l-cry*-/-) worm heads (see **Fig. 5D**), ~2 months-old immature worms were genotyped and separated based on the genotype in different boxes, and starved, one week before sampling (ZT10, new moon [NM] phase). In both experiments, library preparation was carried out using the NEB PolyA kit, to enrich for mRNA. Samples were sequenced on Illumina platform (NovaSeq S4, PE 150). For this experiment, datasets have been analyzed twice independently, and only candidates emerging from both analyses were reported in text and figures (see Table S2 for independent datasets). In the experiment assessing transcriptional profiles of *nr0b1/2* (*nr0b1/2*+/+, *nr0b1/2*+/-, *nr0b1/2*-/-) and *l-cry* (*l-cry*+/+ and *l-cry*-/-) worms (see **Fig. 5A**), 15-34 segments-long immature worms were starved for ~5-7 days, and cut into two equally long parts in the 3 days spanning NM: the most posterior part was used to identify worm´s genotype by PCR and Sanger sequencing, whereas the most anterior one (including the head) was collected in 30 µL of RNAzol, snap-frozen in liquid nitrogen, and stored at -80°C. 3 anterior parts of the same genotype were pooled for each biological replicate and RNA extracted. For the different genotypes, BRs obtained were: *l-cry*+/+ (6 BRs), *l-cry*-/- (6 BRs), *nr0b1/2*+/+ (6 BRs), *nr0b1/2*+/- (6 BRs, 3 $nr0b1/2^{+/\Delta4}$ and 3 $nr0b1/2^{+/\Delta5}$) and *nr0b1/2*-/- (6 BRs, 3 $nr0b1/2^{\Delta4/\Delta5}$ and 3 $nr0b1/2^{\Delta5/\Delta5}$). In this case, library preparation was carried out using SmartSeq3 technology, and samples were sequenced on Illumina platform (NovaSeq S4, PE 150). Finally, in the experiment designed to investigate transcriptional oscillations associated with the lunar cycle **(Fig. 3G)**, *l-cry*+/+ and *l-cry*-/- premature worms with comparable size were collected and decapitated at ZT10 during every phase/week of the artificial lunar cycle (FM, FM+1 week, NM, NM+1 week) for 2 full cycles. Premature animals showing visible/clear signs of sexual metamorphosis were excluded from the analysis. Heads were quickly snap-frozen in liquid nitrogen and stored at -80°C. All animals were starved ~7-10 days before sampling. For each BR (5 BRs for each genotype/timepoint combination), RNA was extracted from 4 premature worm heads using the BioZym kit (Zymo Research).

For the first three RNAseq experiments, raw reads were subjected to quality control using FastQC, and then trimmed using Trimmomatic (v0.39, (*128*)) to remove low-quality bases and adapter sequences. Trimmed reads were then mapped onto both a reference draft genome (*129*) and a reference transcriptome (*130*). Both reference transcriptome and draft genome were automatically annotated using the best hit generated from BLAST searches on Uniprot/Swissprot and RefSeq databases. For data visualization, all hits having no similarity in both databases were excluded (see **table S1-3** for full datasets). For transcriptome mapping and quantification, Salmon (v1.10, (*131*)) was used, while for genome alignment, STAR (2.7.11b, (*132*)) was employed with quantification performed using FeatureCounts (v2.0.3, (*133*)). The resulting count matrices were imported into R (R Core Team (2024). R: A language and environment for statistical computing. R Foundation for Statistical Computing, Vienna, Austria. URL https://www.R-project.org/), and differential expression analysis was carried out using DESeq2 (v1.40.2, (*134*)). Quantitative Venn diagrams were generated using the DeepVenn web tool (https://www.deepvenn.com/).

For the fourth RNAseq experiment (transcriptional oscillations over the lunar cycle), raw reads were subjected to quality control using FastQC, and then trimmed using Trimmomatic (v0.39). Trimmed reads were then aligned to a *Platynereis* reference draft genome (*129*) using STAR (v2.7.11b, (*132*)) and quantified using FeatureCounts (v2.0.6, (*133*)). To identify cyclically expressed transcripts, we used the RAIN (Rhythmicity Analysis Incorporating Nonparametric Methods) algorithm, a method designed to detect rhythmicity in time-series data, accounting for both symmetrical and asymmetrical cycles (*135*). Preprocessed RNAseq read counts were normalized using the DESeq2 package (v3.19) in R (v4.4.1). After normalization, cyclic transcript detection was performed using the RAIN package (v1.0.1). The RAIN algorithm was applied as independent sampling mode, target period of 4 weeks, and peak shape of 0.3, 0.7. P values were then corrected for multiple testing (multiple phases and multiple samples) using the Benjamini-Hochberg method.

<u>qRT-PCR</u>

*l-cry+/+* and *l-cry-/-* immature worms were used to assess the expression levels of *nr0b1/2* in the different artificial lunar phases resulting from standard laboratory LDM. Worms were starved for ~1 week before the sampling, quickly decapitated at ZT10, and heads snap-frozen in liquid nitrogen and stored at -80°C. For each biological replicate, mRNA was extracted from 4-5 worm heads using the BioZym kit (Zymo Research) and reverse transcription carried out using the LunaScript kit (New England Biolabs). qRT-PCR was performed using the Luna Master Mix (New England Biolabs) and the QuantStudio 3 System (Applied Biosystems). mRNA levels were normalized to those of *cdc5* and *rps9* housekeeping genes (*27*), and relative quantification performed by using the $2^{-\Delta\Delta CT}$ method. Results were analyzed using Two-way ANOVA with Sidak´s multiple comparison test: * p<0.05; ** p<0.01; *** p<0.001; **** p<0.0001. Area under the curve (AUC) has been calculated as in (*27*): * p<0.05; ** p<0.01; *** p<0.001; **** p<0.0001. Statistical analysis was performed using Graph Pad Prism v. 9. Primers used were: *nr0b1/2* F: 5'- TCCATCCAGAACTTCATCAGG-3', R: 5'- CGAACTCTCCATTCTCAAGTCC -3'; *cdc5* F: 5'-CCTATTGACATGGACGAAGAT G-3', R: 5'-TTCCCTGTGTGTTCGCAAG-3'; *rps9* F: 5'-CGCCAGAGAGTTGCTGACT-3', R: 5'-ACTCCAATACGGACCAGACG-3'.

<u>Combined *in situ* Hybridization Chain Reaction (HCR) and Immunohistochemistry (IHC)</u>

Simultaneous HCR and IHC labeling was performed according to (*136*). Briefly, worm heads were sampled after being exposed to constant darkness (DD) at circadian time (CT) 6 and fixed in 4% paraformaldehyde (PFA) for 1 hour at room temperature. We chose this timepoint as both artificial and naturalistic sunlight were shown to promote L-Cry degradation (*22*, *27*). Samples were dehydrated using increasing concentrations of methanol (MeOH) and finally stored in 100% MeOH at -20°C. Then, samples were first rehydrated and digested for 5 minutes using Proteinase K, rinsed twice in Glycine, each time for 1 minute, washed twice in 1x PTW (1x PBS/ 0,1% TWEEN20), post-fixed in 4% PFA for 20 min, and finally washed 3 times for 5 minutes each (all steps performed on ice). Following an initial equilibration and pre-hybridization of worm heads in probe hybridization buffer at 37°C (for 1 hour), samples were incubated overnight in probe solution (specific for the detection of *nr0b1/2*) at 37°C. The next day, several washes were performed using pre-heated probe wash buffer at 37°C, followed by 2 washes of 5 minutes in 5x SSCT (Saline-sodium citrate buffer with 0,1% TWEEN20) at room temperature. For the detection of the HCR probe, hairpins compatible with the probe amplifier were prepared and mixed with the amplification buffer. At this stage, anti-*Pdu*-L-Cry antibody was added (1:100) to the hairpin-mix, and samples were incubated overnight. Then, once amplification terminated (day after), the primary antibody was washed off using 5x SSCT at room temperature (~70 minutes in total), followed by 2 washes of 5 minutes in 1x PTW. Samples were then incubated overnight at 4°C with the secondary antibody (AF555 goat anti-mouse, 1:500). 2 final washes of 15 minutes in 1x PTW were performed, and samples mounted using SlowFade Diamond Antifade Mountant (Invitrogen). Imaging was performed using a Zeiss LSM700 inverse confocal microscope with LD LCI Plan-Apochromat 25x/0.8, Imm. Corr. DIC M27, Plan-Apochromat 40x/1.3 Oil DIC M27 lenses. Microscope parameters were set individually for every image, and using Fiji (Image J, (*127*)) the contrast (linear adjustment) and brightness were enhanced.

Generation and genotyping of *nr0b1/2* mutant worms

TALENs design and preparation were carried out as in (*137*). We targeted exon 1 (~1.3 kb) in the *nr0b1/2* locus, and the TALENs recognition site was: 5'-GGGAGTGTTTCATCAcccagtgatagccagGCTGCTGCTGCTGCT-3'. *nr0b1/2+/-* (*nr0b1/2*$^{+/\Delta5}$ or *nr0b1/2*$^{+/\Delta4}$) and *nr0b1/2-/-* (*nr0b1/2*$^{\Delta5/\Delta5}$ or *nr0b1/2*$^{\Delta4/\Delta5}$) strains were generated from single founders, carrying a 5-bp or a 4-bp deletion in the *nr0b1/2* coding region (**Fig. 4A**). These deletions created a frameshift that introduced premature stop codons after 7 and 21 amino acids, respectively (**Fig. 4A**). To identify worm genotypes, the region encompassing TALENs-targeted site was amplified (primers F: 5'-AAACGTTTAAAAGGATTGTACCCGG-3', R: 5'-CCGACGATTTGGGCGCTTCC-3'), and the restriction digestion performed using *BtsIMutI* (New England Biolabs). Alternatively, following PCR amplification, amplicons were sequenced via Sanger sequencing using forward primers, and sequences obtained compared bioinformatically using CLC Main Workbench (Qiagen). Notably, the generation of *nr0b1/2*$^{\Delta5/\Delta5}$ homozygous and *nr0b1/2*$^{\Delta4/\Delta5}$ trans-heterozygous mutants was possible only by isolating worms early in development in individual wells. Indeed, *nr0b1/2-/-* worms appeared to suffer from fitness disadvantage under competitive conditions (presence of the other genotypes, or simply other individuals, within the same space), and they never reached maturation in the presence of *nr0b1/2+/+* or *nr0b1/2+/-* animals (**Fig. 4B**). Specifically, to obtain *nr0b1/2-/-* homozygous worms, 95 ~10-30 segments-long individuals were randomly selected from several age-matched heterozygous mating. Worms were genotyped by amputating their posterior half, before raising them for their entire life in isolated wells.

Phylogenetic study and sequence analysis

To reconstruct phylogenetic relationships for the major candidates identified in RNAseq experiments (*nr0b1/2*, *dhr11*, *cp17a1/2*, *cp46a1*, *pdp1*, *cwo*, *nr5a1/2*, *errα/β/γ*, *s5a1-like*, *burs-like*, *dmrt1*, *dmrt3/c2*, other *nuclear receptors*, *ovol1/2/3*), *Platynereis* transcripts (generated mapping reads on both a cDNA database with genome-predicted transcripts (*129*), and a previously published transcriptome, (*130*)) were used to identify the putative coding sequences and obtain the corresponding translations using ORFfinder (NCBI). For the other species, protein sequences were retrieved by Basic Local Alignment Search Tool (BLAST) from NCBI databases. Hit sequences were aligned using MUSCLE (*138*). Maximum likelihood phylogenies were generated using the IQ-TREE (*139*) web server, with default settings. Consensus trees generated were visualized using the Interactive Tree of Life tool (*140*), and rooted using a protein outgroup (outside the protein group of interest) identified by BLAST searches (*30*). Protein sequences used for phylogenetic reconstructions and their respective accession numbers can be found in **table S7**. Sequences of *Platynereis* transcripts identified and phylogenetically validated as described above are available under the following GenBank identifiers: *nr0b1/2* (…), *dhr11* (…), *cp17a1/2* (…), *cp46a1* (…), *pdp1*

(…), *cwo* (…), *nr5a1/2* (…), *errα/β/γ* (…), *s5a1-like* (…), *burs-like* (…), *dmrt1* (…), *dmrt3/c2* (…), other *nuclear receptors* (…), *ovol1/2/3* (…).

For the analysis of Nr0b1/2 domains and motifs conservation across bilaterian species, protein sequences obtained as described above were aligned in CLC Main Workbench v. 22, and boundaries of ligand- and DNA-binding domains displayed based on human DAX-1 (see (*141*)). For the identification and visualization of nuclear receptor (NR)-binding box motifs (typical of NRs coactivators and corepressors), we searched for and displayed all LXXLL and LXXXL (LXXLL variants with the leucine at position 4 substituted) (overall called LXXLL-related) consecutive residues (see (*86*)), including those overlapping, as several are evolutionary conserved (see **fig. S11**). Protein sequences used to generate the alignment and their respective accession numbers can be found in **table S7**.

### Supplementary Text

#### Details to *Platynereis* post-larval development

In brief, after larval settling, immature worms grow progressively adding new segments posteriorly. Afterwards, in coordination with body growth at approximately the size of 30-40 segments, germline precursors start proliferating (*37*), and then differentiating into spermatogonia or oocytes in male and female worms, respectively. Germline differentiation in the two sexes represents the hallmark of worms entering the premature stage. Finally, after a period of continuous growth by segments addition, which accompanies spermatogonia and oocytes development, premature worms enter the last phase of sexual maturation (sexual metamorphosis or epitoky), when secondary sexual characters becomes clearly visible, an event that anticipate by 7-10 days on average the achievement of full sexual maturity (*30*). During sexual metamorphosis, worms undergo significant physiological and morphological changes, which include muscle and parapodia rearrangement from crawling to swimming forms, the reabsorption of the gut (and the consequent cessation of feeding), and a massive eye growth. However, the most notable feature consists of male and female worms turning their immature/premature mimetic coloration into red and yellow liveries (with mature oocytes colour contributing to the second) (ref.*18*, and **Fig. 1A**). Once gametes become mature and worms fully metamorphosed, *P. dumerilii* individuals leave their shelters (i.e. algae) to engage in nuptial dances which terminate with the emission of gametes in the water column (spawning). This represents their unique reproductive event, which is shortly followed by their death (*18*). Interestingly, in nature both sexual metamorphosis and reproduction are timed with specific phases of the lunar cycle to increase the probability of gametes encounter (*18*), a synchronization that can be mimicked also in laboratory conditions (*22*, *27*, *30*).

#### Definition of peptidergic GPCRs

Among the many candidates obtained from RNAseq experiments, we identified several *G protein-coupled receptors* (*gpcrs*). As our attempts to unambiguously reconstruct the phylogenetic relationships of some of them were unsuccessful, we named them according to our initial automated annotation based on BLAST searches using Uniprot/Swissprot and RefSeq databases or, in two cases as *gpcrx* and *gpcry*, respectively. The latter choice was dictated by the fact that additional BLAST searches and homology analyses indicated that the two initial annotations appeared incorrect. Indeed, blasting the single mapped read annotated as *Green-sensitive opsin* and two (out of seven) of those annotated as *Rhodopsin*, resulted in non-opsin receptors, primarily peptidergic, as best hits (see **table S5** and **table S6**, respectively). This is in part due to the fact that blasting the identified non-opsin GPCRs

against vertebrate datasets resulted in opsin-type GPCRs as best hits. Consistently, using insect and mollusc databases, reciprocal best BLAST hits were primarily peptidergic GPCRs. Specifically, *gpcry* reported in the Venn diagram of **Fig. 5H** and **fig. S4F** as co-regulated in both *l-cry* and *nr0b1/2* mutants results as a putative peptidergic receptor (see **table S4** and **table S6**), whereas *gpcry* in the Venn diagram from **fig. S4G** likely represents a *rhodopsin* (see **table S4** and **table S6**).
In summary, we termed some of the identified GPCRs as "peptidergic" based on either of two criteria: 1) the absence of the typical Lysine (Lys, K, in position 296 in the human Rhodopsin) necessary to form a Schiff base linkage when reacting with the cofactor 11-*cis*-retinal, a key mechanism in the context of photoreception (*142*) (see **fig. S22**); 2) the hits resulting from the automated annotation or reciprocal BLAST searches in well studied protostome groups (insects and molluscs, see **table S1-3**, **S5** and **S6**).
Particularly, for *gpcrx*, the most consistently regulated GPCR throughout the study, we provide an alignment including human Rhodopsin and some of the best hits generated by blasting this mapped read against vertebrate, insect, and mollusc databases, to show the absence of the Lys necessary to form the Schiff base (**fig. S22**). All BLAST searches against vertebrate, insect and mollusc genomes are reported in **table S5** and **table S6**.

**Supplementary Figures**

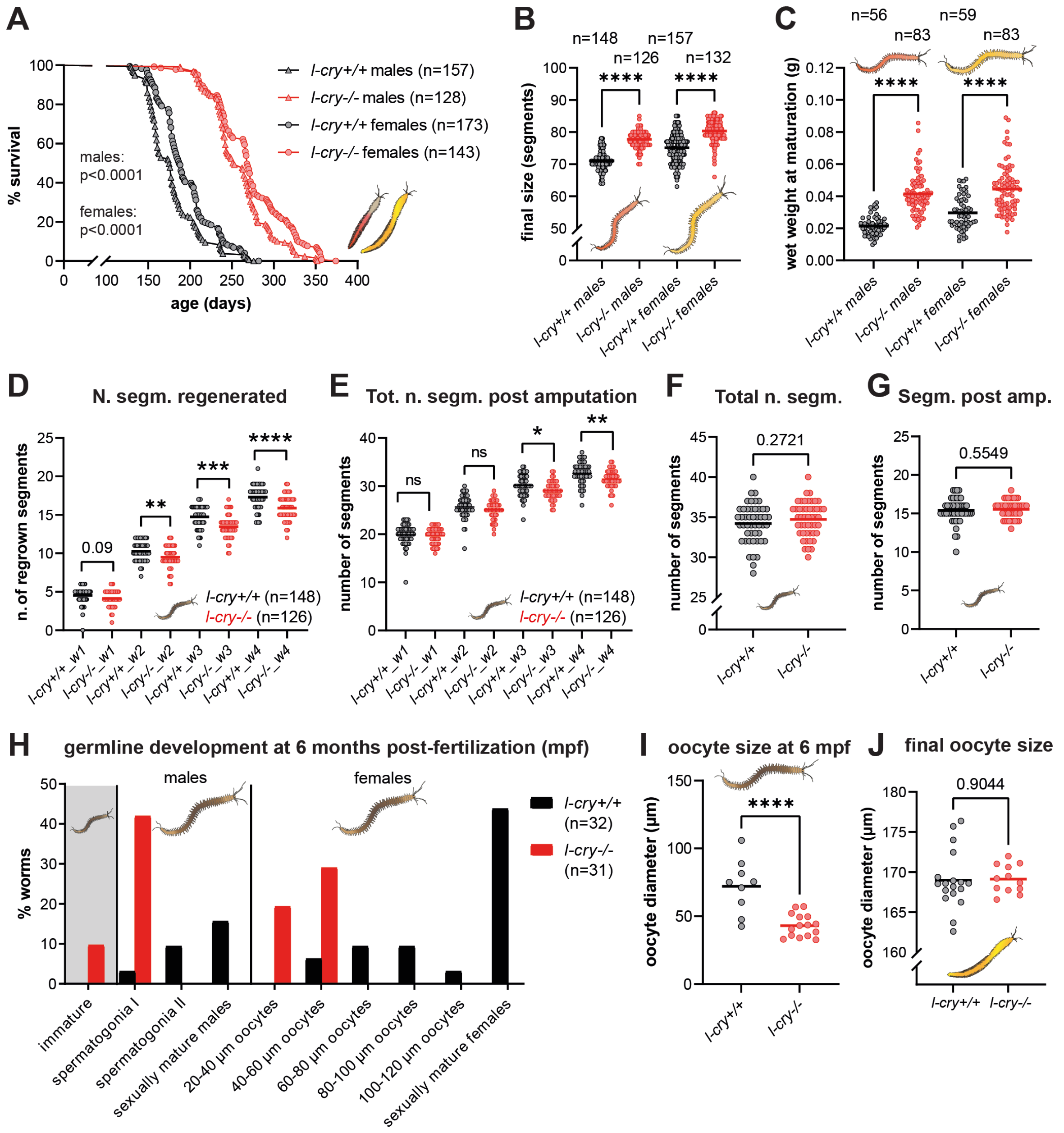


**figure S1: *l-cry-/-* knockouts extend lifespan and final size, delay growth/regeneration and germline development.** **(A)** Survival (maturation) curves of *l-cry+/+* and *l-cry-/-* male and female worms indicating the proportion of individuals that have yet to attain sexual maturity, and thus are still alive (% survival), over time. Lifespan was considered the time-span between fertilization and reproduction/death. N ≥ 128 for each sex/genotype combination. Log-rank (Mantel-Cox) test, $p < 0.0001$ for comparison between genotypes under indicated condition. **(B)** Final size of *l-cry+/+* and *l-cry-/-* male and female worms quantified as total number of segments at maturation. N ≥ 126 for each sex/genotype combination. One-way ANOVA, **** $p < 0.0001$. **(C)** Wet weight of *l-cry+/+* and *l-cry-/-* male and female worms during sexual metamorphosis. N ≥ 56 for each sex/genotype combination. One-way ANOVA, **** $p < 0.0001$. **(D)** Growth rate of age-matched *l-cry+/+* and *l-cry-/-* worms quantified by assessing the number of segments regrown after the amputation of the posterior half of the worm body. w1, w2, w3, w4 = 1 week, 2 weeks, 3 weeks, 4 weeks post amputation. N ≥ 126 for each genotype. T test, ** $p < 0.01$; *** $p < 0.001$; **** $p < 0.0001$. **(E)** Growth rate of age-matched *l-cry+/+* and *l-cry-/-* worms quantified by assessing the number of total body segments after the amputation of the posterior half of the worm body. w1, w2, w3, w4 = 1 week, 2 weeks, 3 weeks, 4 weeks post amputation. N ≥ 126 for each genotype. T test, * $p < 0.05$; ** $p < 0.01$. **(F)** Total number of body segments in the animals used in (D) and (E) prior amputation. T test. **(G)** Total number of body segments in the animals used in (D) and (E) following amputation. T test. **(H)** Characterization of germline development in age-matched *l-cry+/+* and *l-cry-/-* worms after 6 months from fertilization. Bars represent the proportion of wt or knockout worms found at each stage of germline development considered. N ≥ 31 for each genotype. **(I)** Maximum oocytes size (diameter) at 6 months from fertilization in individual *l-cry+/+* and *l-cry-/-* premature females. Oocytes from fully mature females (only present for wt) were excluded. N ≥ 9 for each genotype. T test, **** p < 0.0001. **(J)** Maximum oocytes size (diameter) at reproduction (spawning) in *l-cry+/+* and *l-cry-/-* females. N ≥ 12 for each genotype. T test.

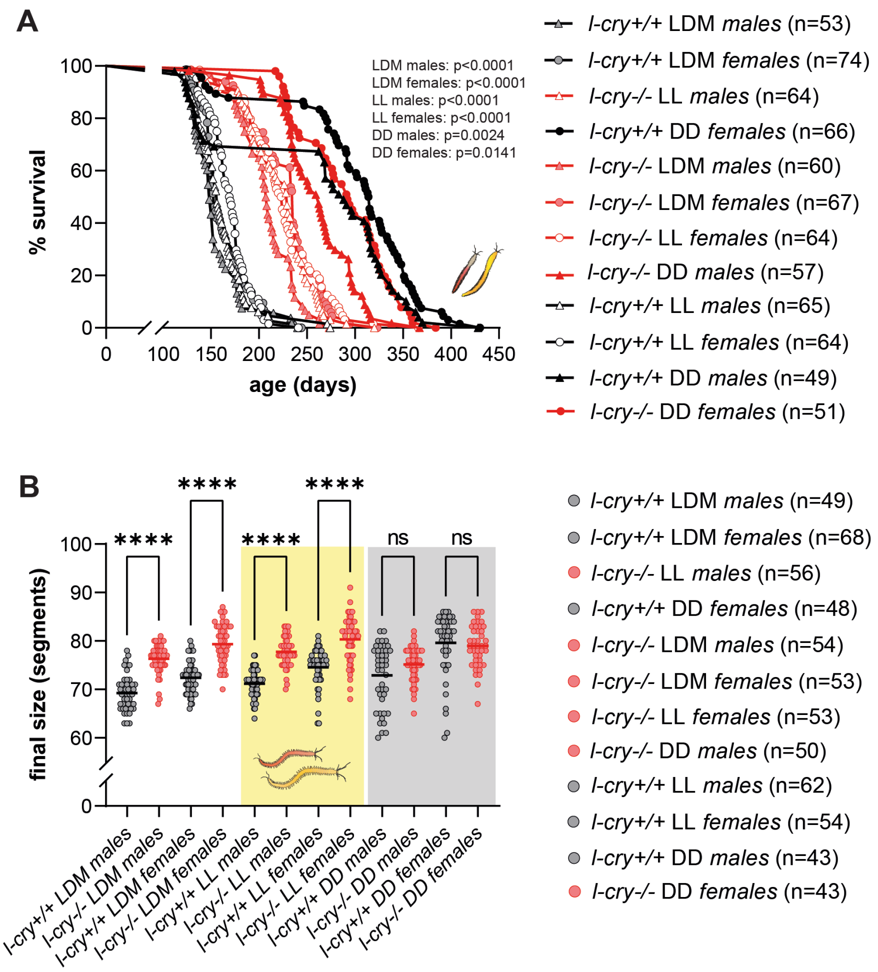


**figure S2: Lifespan and growth extension are L-Cry-and light-dependent in male and female *Platynereis* worms. (A)** Survival (maturation) curves of *l-cry+/+* and *l-cry-/-* male and female worms indicating the proportion of individuals that have yet to attain sexual maturity, and thus are still alive (% survival), over time. Lifespan was considered the time-span between fertilization and reproduction/death. N ≥ 49 for each sex/genotype combination. Log-rank (Mantel-Cox) test, p < 0.0001 refer to comparisons of wt vs. mutants at mentioned condition . **(B)** Final size of *l-cry+/+* and *l-cry-/-* male and female worms quantified as total number of segments at maturation. White background = light/dark (LD 16:8) + 8 nights of naturalistic moonlight; yellow background = light/light (LL); grey background = dark/dark (DD). N ≥ 43 for each sex/genotype combination. T test, **** p < 0.0001.

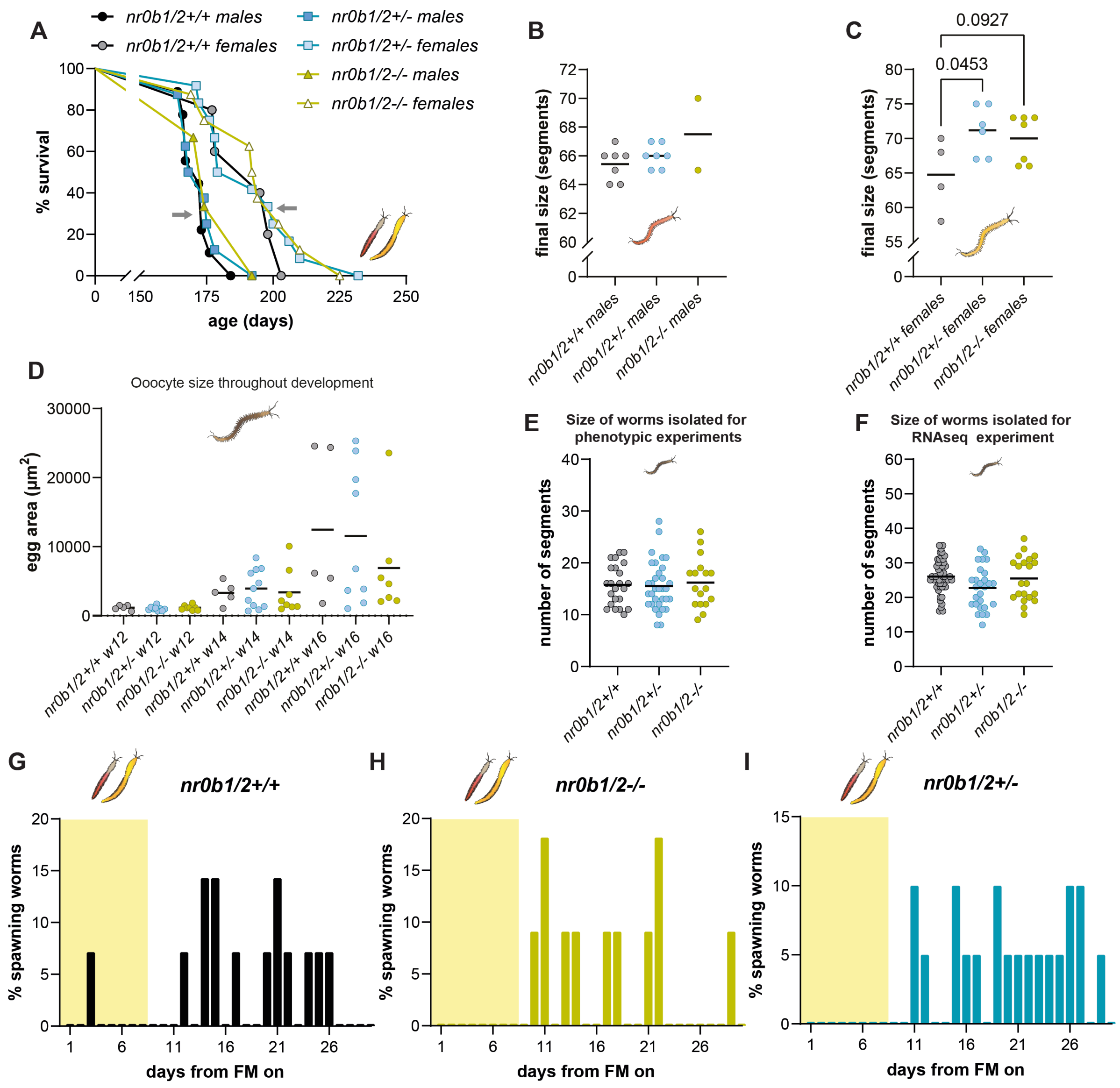


**figure S3: *nr0b1/2* mutant worms partially phenocopy the lifespan and growth extension of *l-cry-/-* mutants, but show no altered lunar rhythms. (A)** Survival (maturation) curves of *nr0b1/2+/+*, *nr0b1/2*+/- and *nr0b1/2*-/- worms indicating the proportion of individuals that have yet to attain sexual maturity, and thus are still alive (% survival), over time. N ≥ 11 for each genotype. **(B)** Final size of *nr0b1/2+/+*, *nr0b1/2*+/- and *nr0b1/2*-/- male worms quantified as total number of segments at maturation. N ≥ 2 for each sex/genotype combination. One-way ANOVA. **(C)** Final size of *nr0b1/2+/+*, *nr0b1/2*+/- and *nr0b1/2*-/- female worms quantified as total number of segments at maturation. N ≥ 4 for each sex/genotype combination. One-way ANOVA. **(D)** Maximum oocytes size (egg area) after 12, 14, and 16 weeks from tail amputation (necessary for genotyping) of *nr0b1/2+/+*, *nr0b1/2*+/- and *nr0b1/2*-/- premature females. N ≥ 5 for each genotype/timepoint. T test. **(E)** Body size of age-matched *nr0b1/2+/+*, *nr0b1/2*+/- and *nr0b1/2*-/- immature worms (with some successfully surviving till sexual maturation and used in Fig. 4C, D, E and fig. S3A-C, G-I) individually isolated early in development and before tail amputation (necessary for genotyping), assessed by counting segments number. N ≥ 17 for each genotype. One-way ANOVA. **(F)** Body size of age-matched *nr0b1/2+/+*, *nr0b1/2*+/- and *nr0b1/2*-/- immature worms whose head was used for RNAseq (see Fig. 5A, C and fig. S4A, C, D), before tail amputation for genotyping, assessed by counting segments number. N ≥ 21 for each genotype. One-way ANOVA. **(G-I)** Reproductive (spawning) frequencies (expressed as %) of *nr0b1/2+/+* (G), *nr0b1/2*-/- (H), and *nr0b1/2*+/- (I) male and female worms over the lunar month in the lab (with 8 nights of artificial moonlight, represented by yellow rectangles). N ≥ 14 for each genotype.

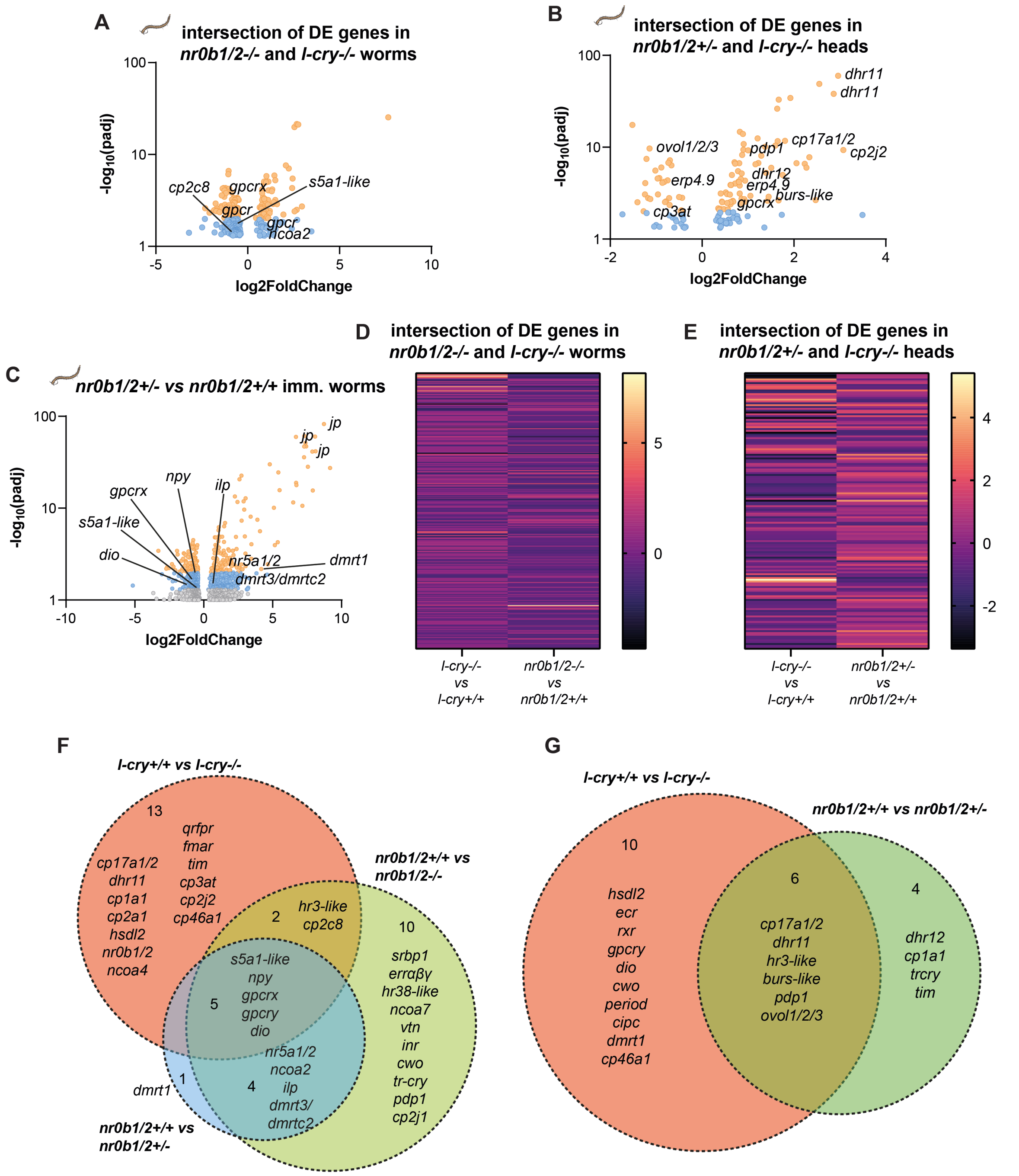


**figure S4: *nr0b1/2* regulates endocrine pathways and transcriptional regulators associated to sexual development/reproduction and circadian rhythms, partially overlapping with L-Cry targets. (A, B)** Volcano plot visualizing only the transcripts regulated in both *nr0b1/2+/+* vs *nr0b1/2-/-* and *l-cry+/+* vs *l-cry-/-* worms (A), and *nr0b1/2+/+* vs *nr0b1/2+/-* and *l-cry+/+* vs *l-cry-/-* worm heads (B) comparisons, respectively, with a particular emphasis on the endocrine system. To display the relationship between adjusted p value (padj) and Fold Change, these values have been converted to $-\log_{10}$(padj) and log2FoldChange, respectively. For the sake of clarity, in volcano plots, only transcripts with padj $\leq 0.1$ are displayed. Orange dots: padj $\leq 0.01$; blue dots: padj $\leq 0.05$. **(C)** Volcano plots summarizing major expression changes between *nr0b1/2+/+* and *nr0b1/2+/-* immature worms, focusing primarily on the endocrine system and actors involved in sexual development. **(D, E)** Heat maps displaying similarities and differences in the molecular signatures deriving from *nr0b1/2+/+* vs *nr0b1/2-/-* and *l-cry+/+* vs *l-cry-/-* worms (D), and *nr0b1/2+/+* vs *nr0b1/2+/-* and *l-cry+/+* vs *l-cry-/-* worm heads (E) comparisons, respectively. Every horizontal line corresponds to the fold change of the same transcript in both analyses. **(F, G)** Quantitative Venn diagrams showing transcripts significantly and jointly regulated in *l-cry+/+* vs *l-cry-/-*, *nr0b1/2+/+* vs *nr0b1/2-/-*, and *nr0b1/2+/+* vs *nr0b1/2+/-* worms (F), or in *l-cry+/+* vs *l-cry-/-*, and *nr0b1/2+/+* vs *nr0b1/2+/-* worm heads (G) comparisons, respectively, focusing primarily on those related to endocrine pathways, sexual development/reproduction, and biological rhythms.

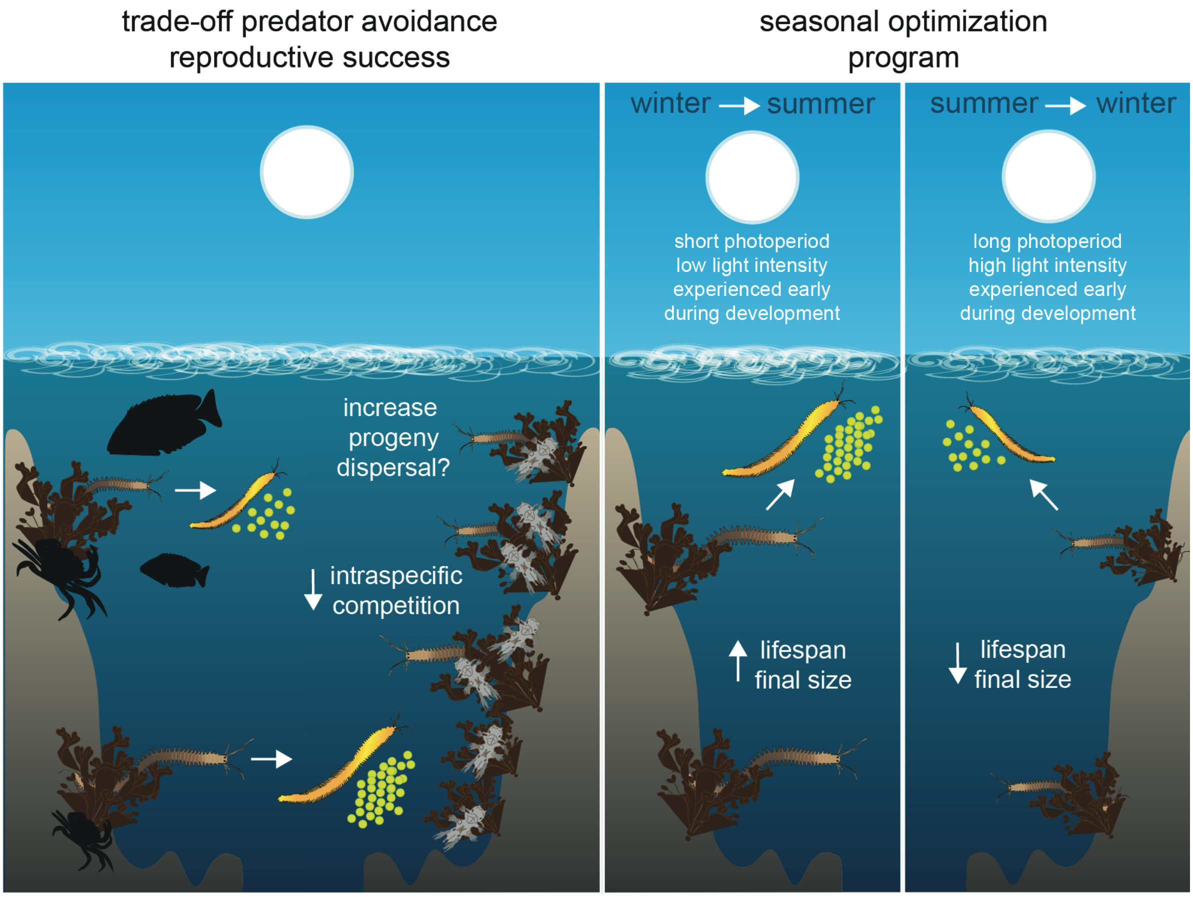


**figure S5:** Ecological model of the hypothesized adaptive value of light-dependent lifespan and growth extension.

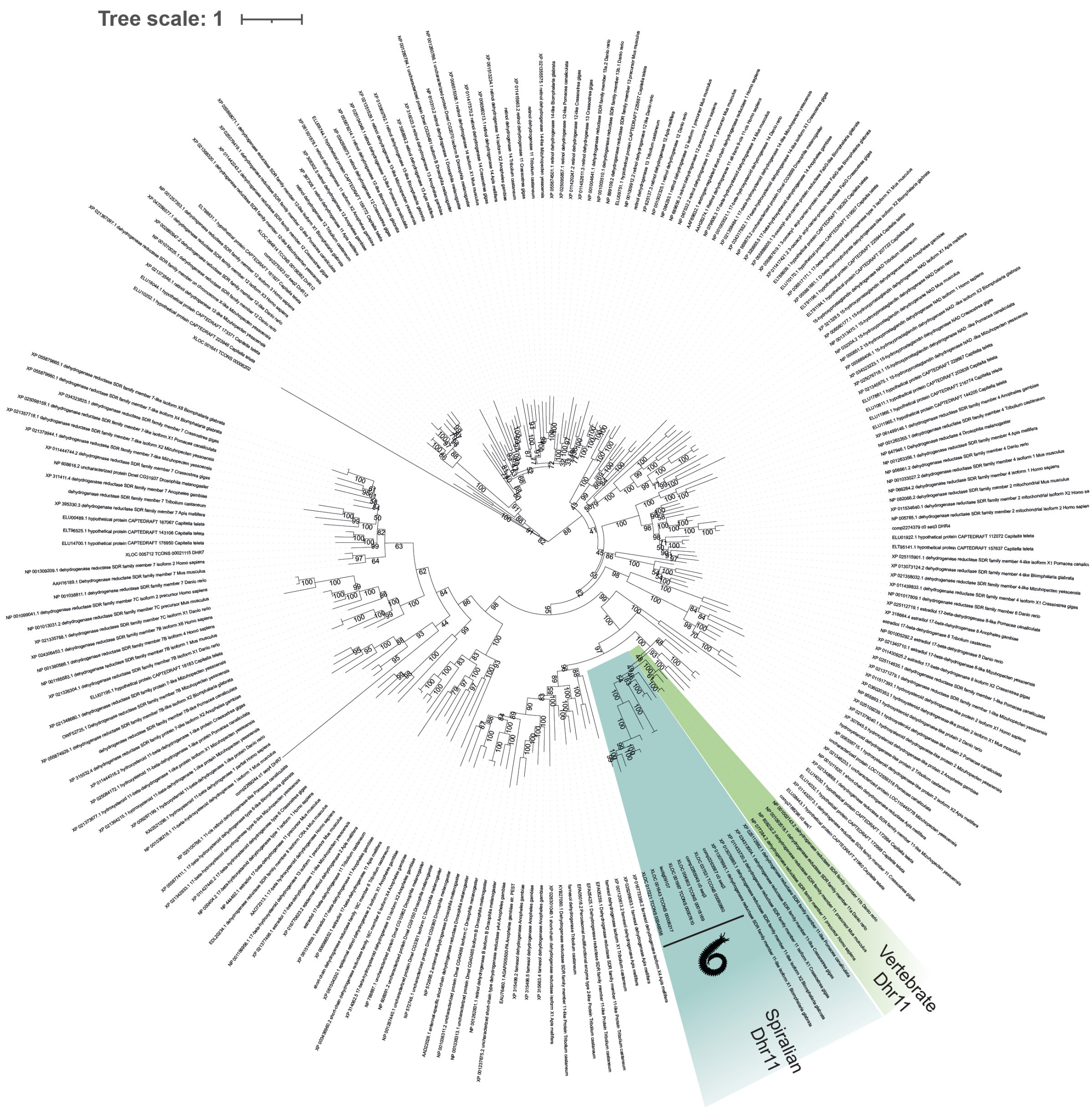


**figure S6:** Phylogenetic tree reconstruction of bilaterian Dhr11 proteins.

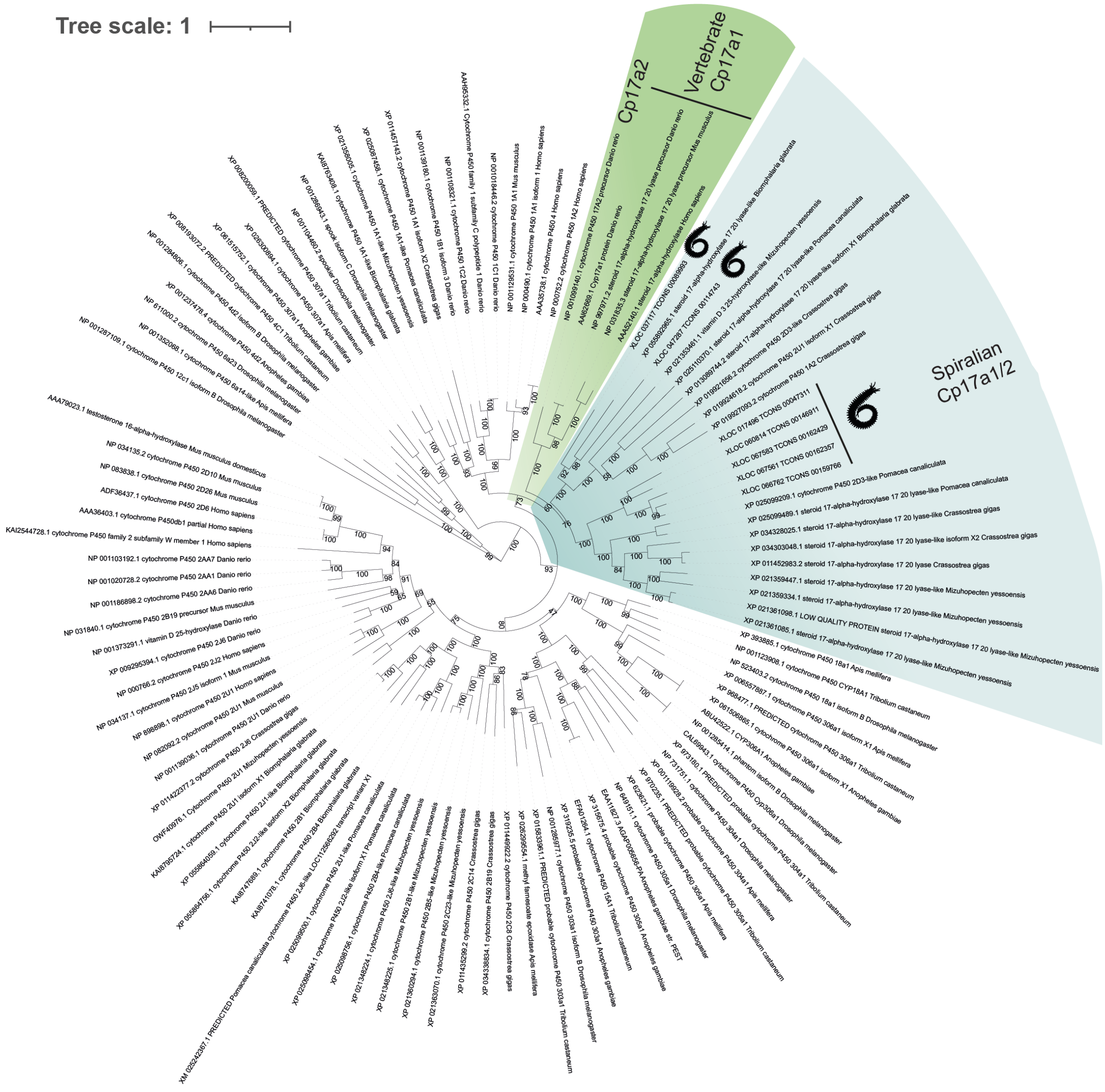


**figure S7:** Phylogenetic tree reconstruction of bilaterian Cp17a1/2 proteins.

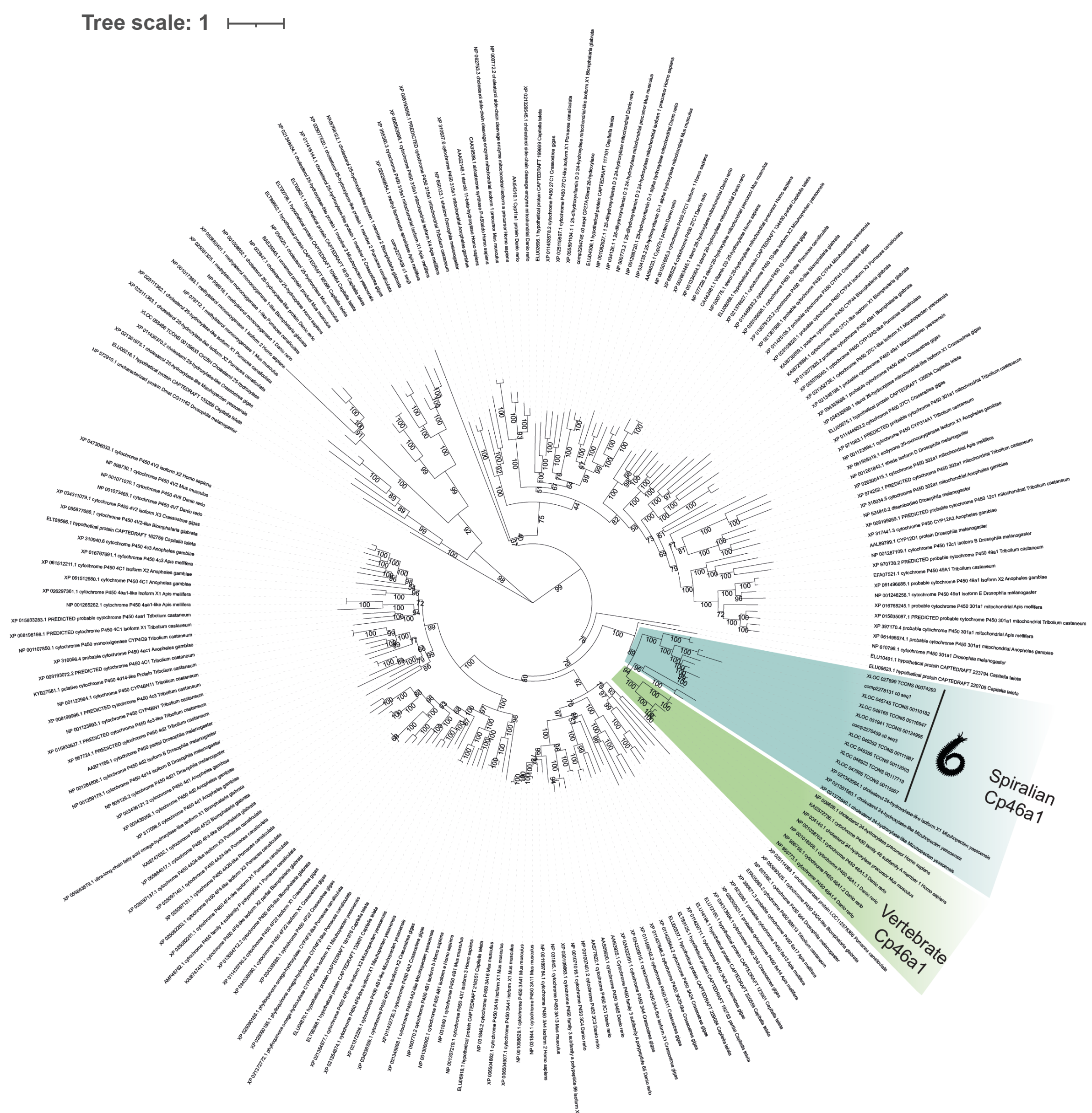


**figure S8:** Phylogenetic tree reconstruction of bilaterian Cp46a1 proteins.

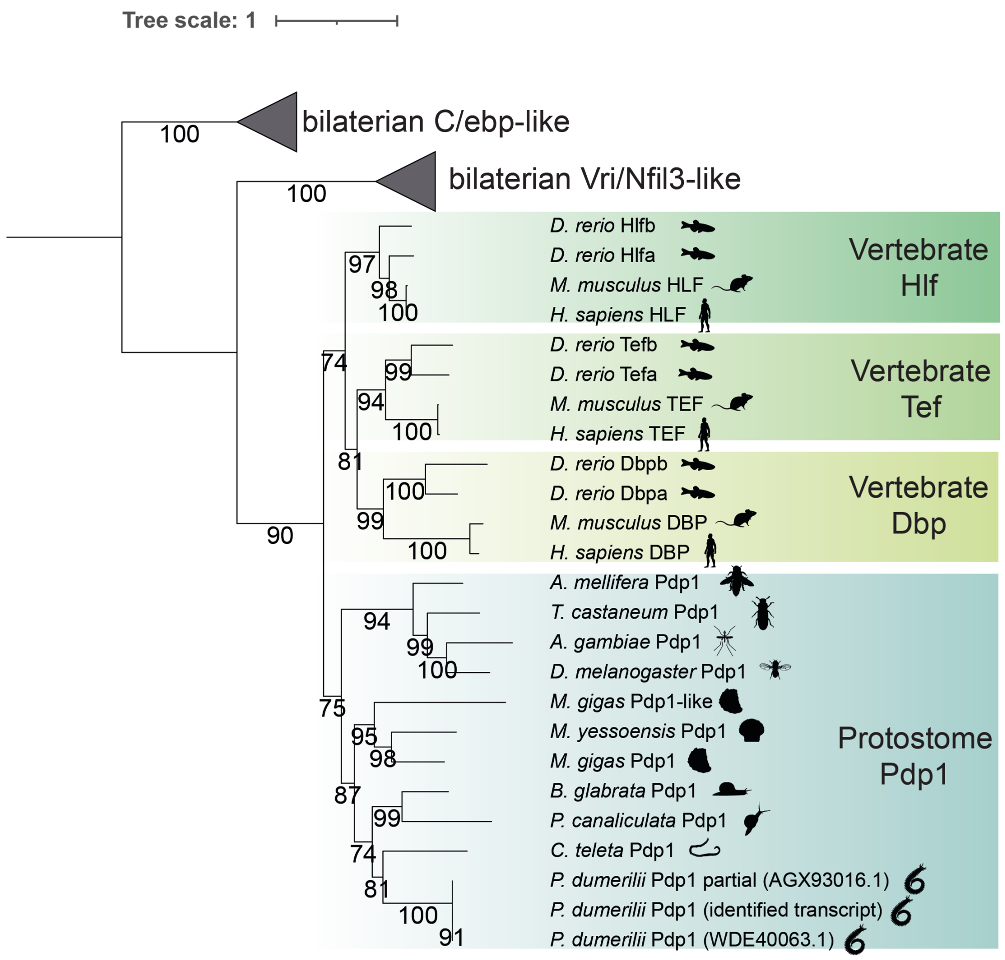


**figure S9:** Phylogenetic tree reconstruction of bilaterian Pdp1, Hlf, Dbp and Tef bZIP transcription factors.

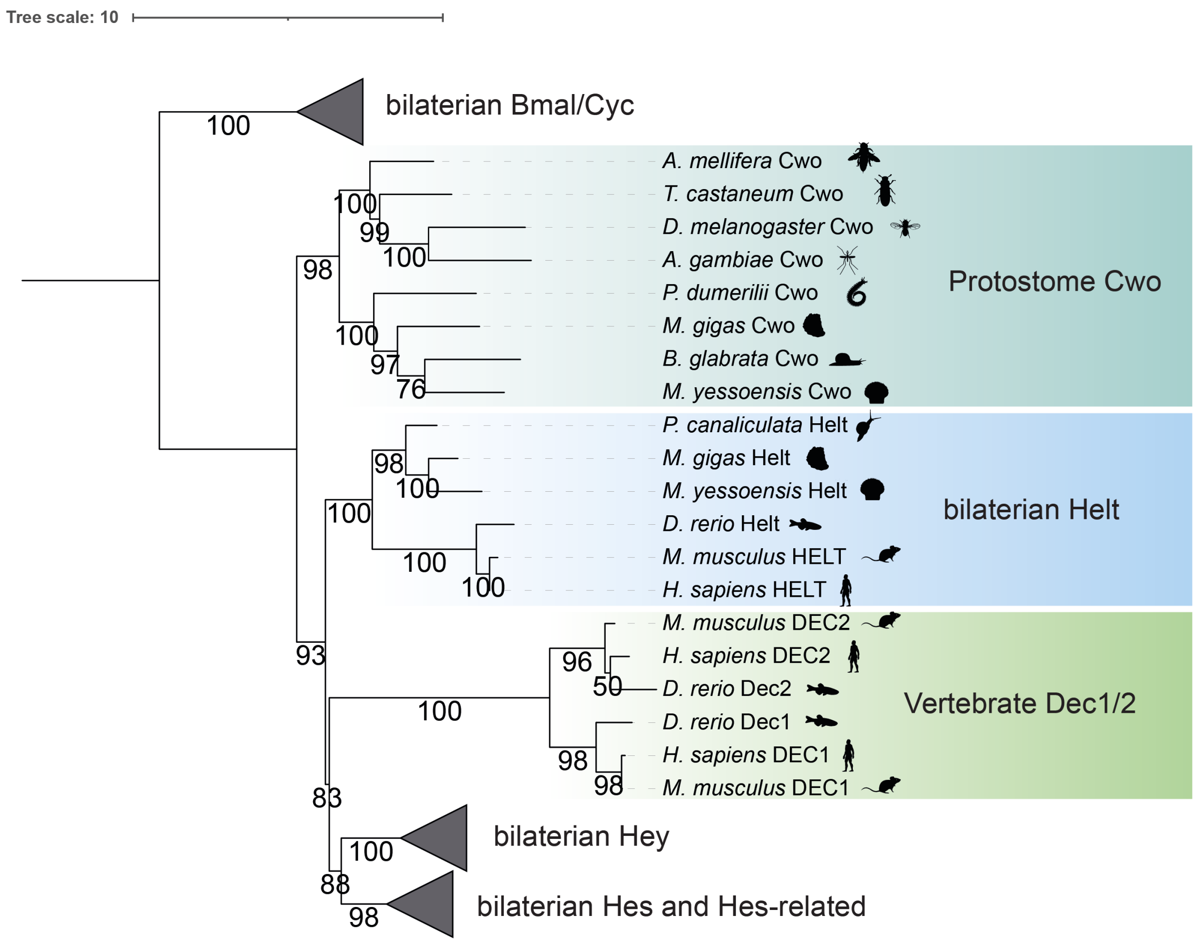


**figure S10:** Phylogenetic tree reconstruction of bilaterian Cwo, Dec1/2, and closely-related transcriptional regulators.

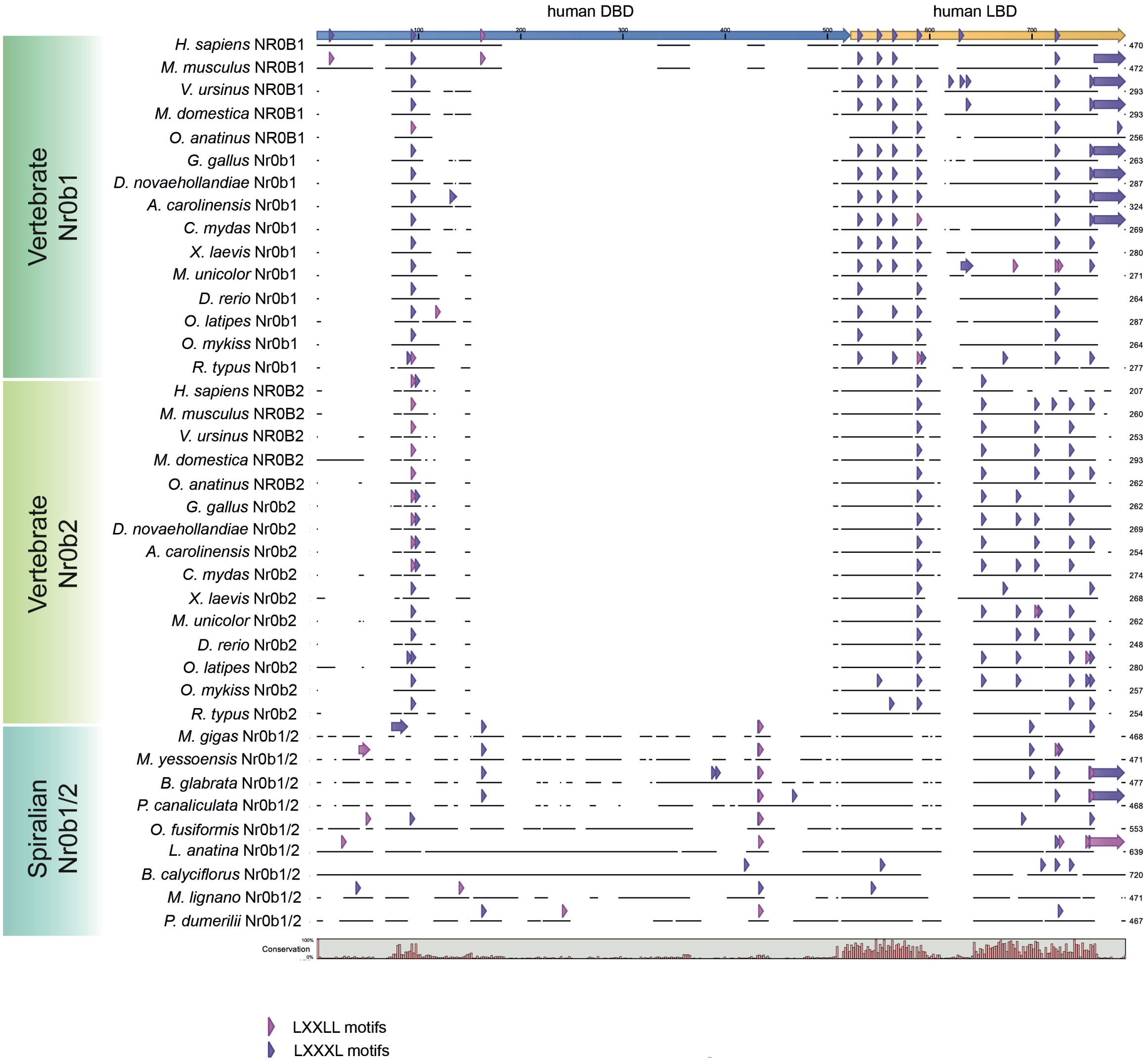


**figure S11:** Alignment of bilaterian Nr0b1/2 nuclear receptors from representative vertebrate and spiralian species.

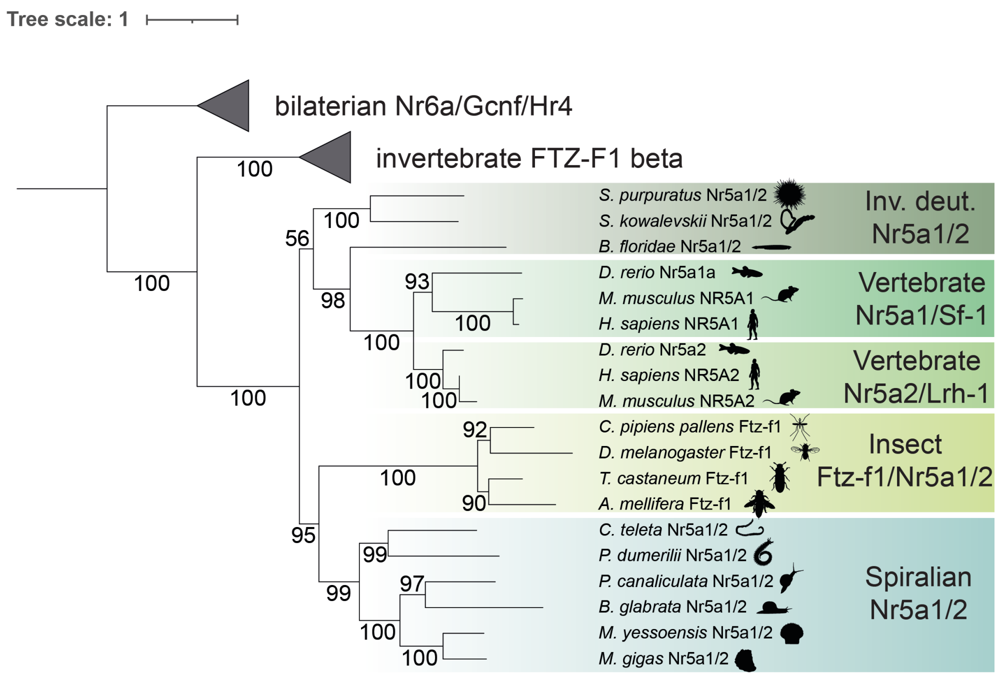


**figure S12:** Phylogenetic tree reconstruction of bilaterian Nr5a1/2 nuclear receptors.

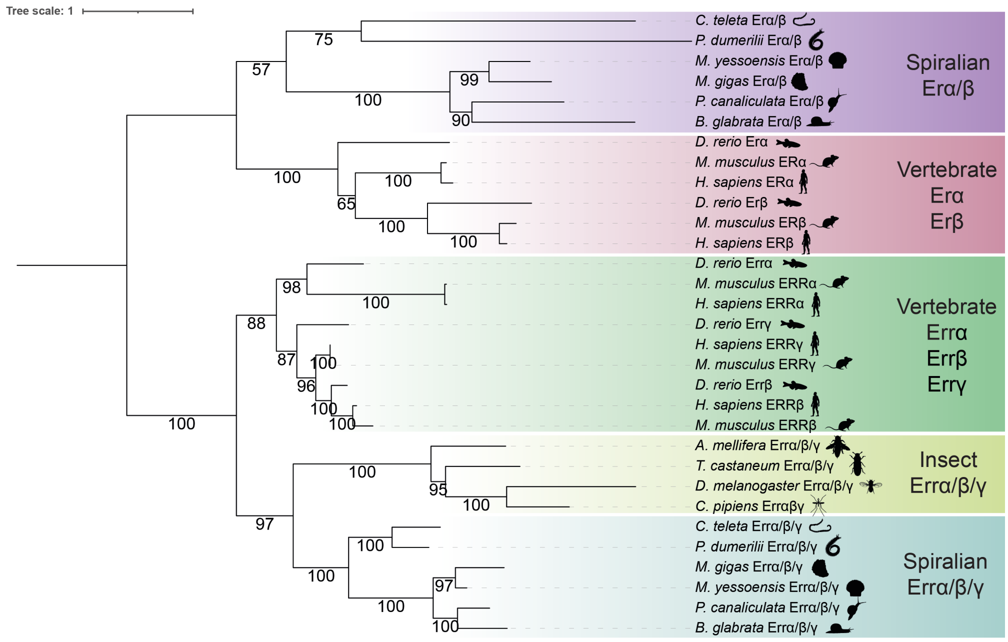


**figure S13:** Phylogenetic tree reconstruction of bilaterian Estrogen (Er) α/β and Estrogen-related (Err) α/β/γ nuclear receptors.

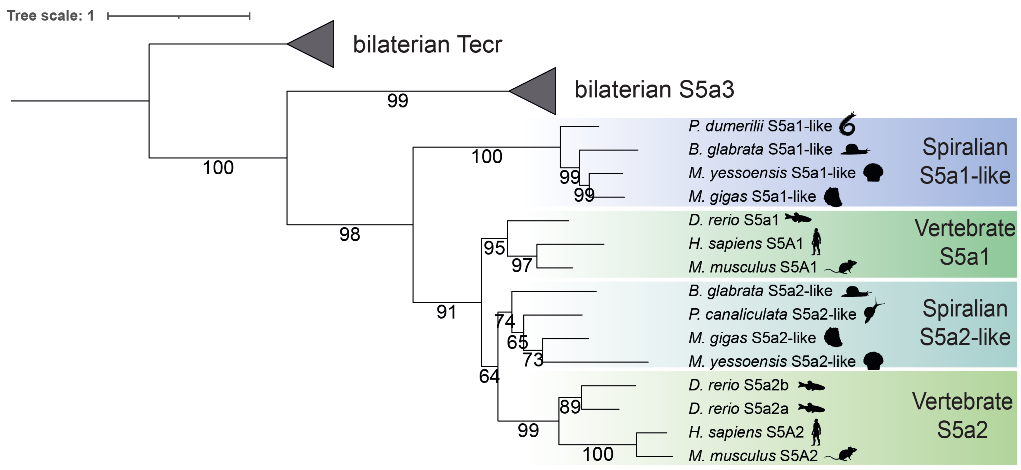


**figure S14:** Phylogenetic tree reconstruction of bilaterian steroid 5-alpha (S5a) reductase-like enzymes.

Fig. S14

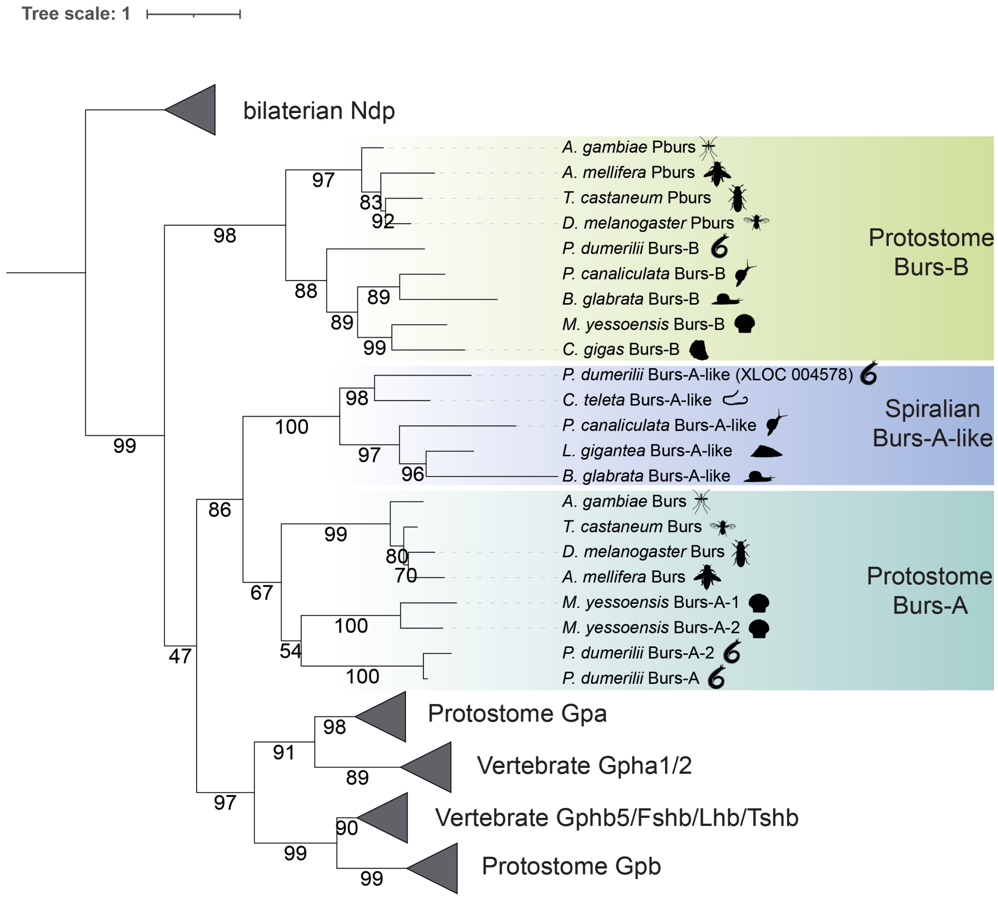


**figure S15:** Phylogenetic tree reconstruction of the bilaterian bursicon neuropeptide family of glycoprotein hormones.

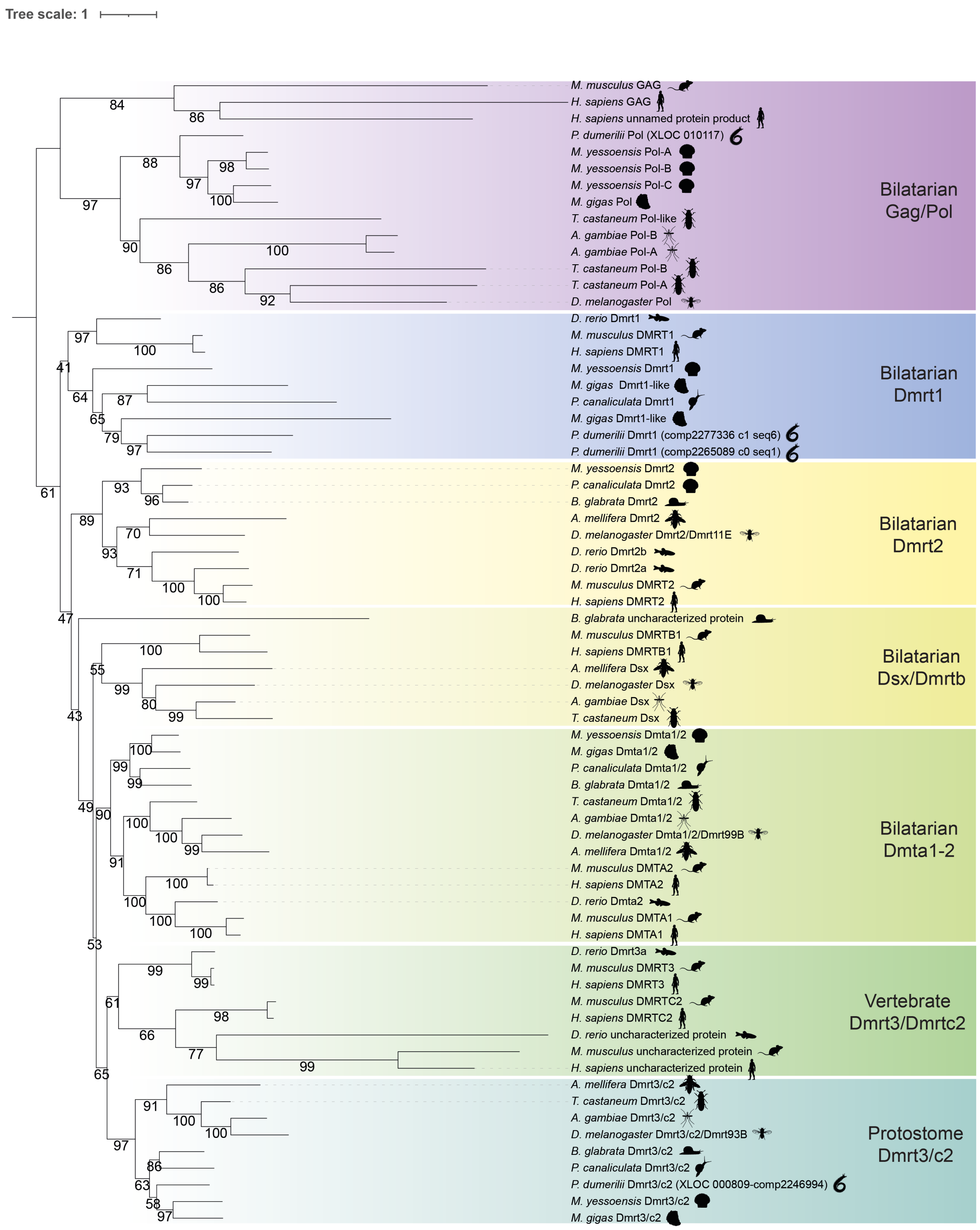


**figure S16:** Phylogenetic tree reconstruction of bilaterian Doublesex and mab-3 related transcription factors 1 (Dmrt1) and **Doublesex- and mab-3-related transcription factors 3/c2(Dmrt3/c2)** proteins.

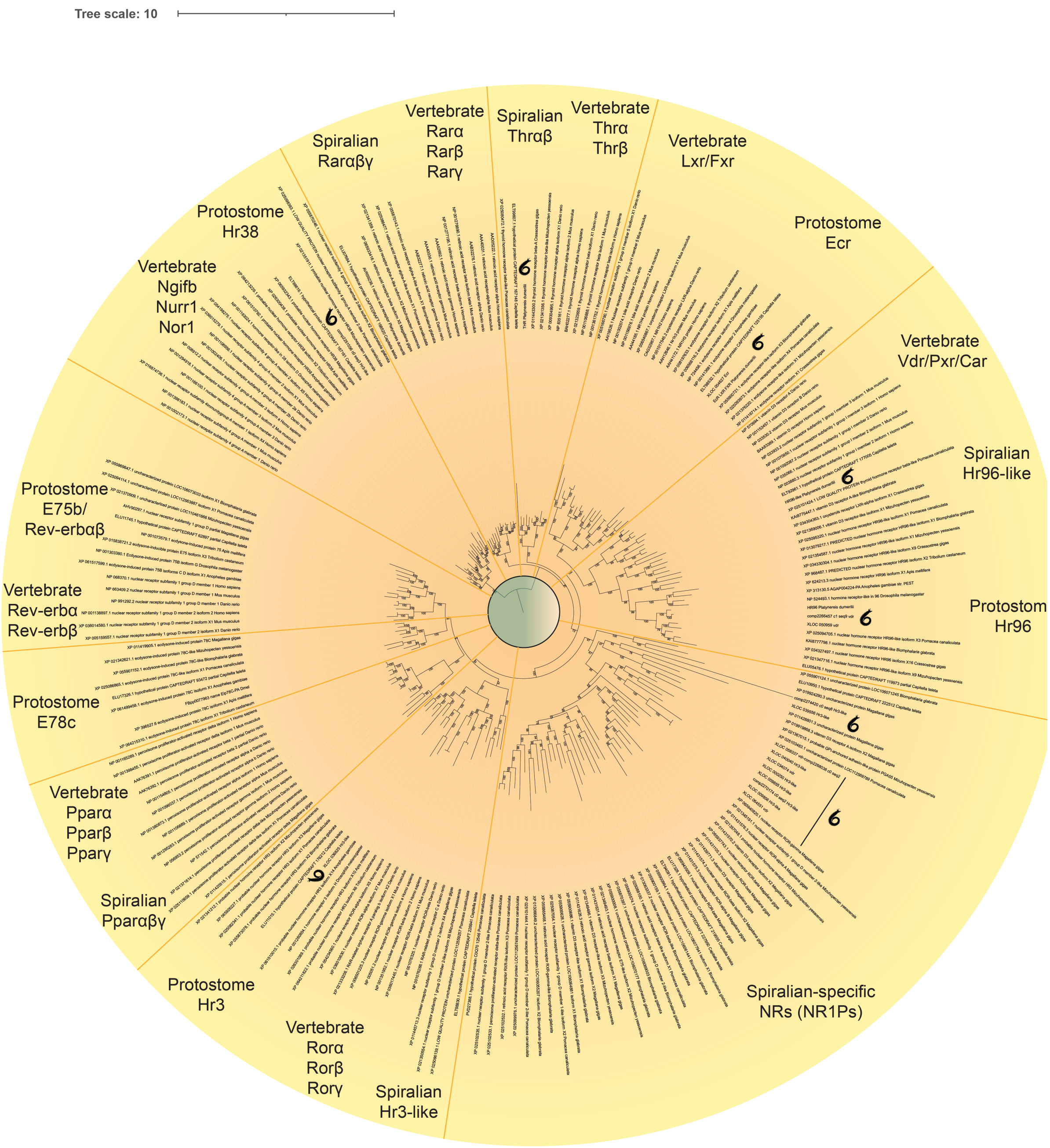


**figure S17:** Phylogenetic tree reconstruction of bilaterian Rar, Thr, Lxr/Fxr, Ecr, Vdr/Pxr/Car, Hr-96, Hr96-like, Hr3, Hr3-like, Ror, Ppar, E78c, E75b/Rev-erb, Ngifb/Nurr1/Nor1, Hr38, and Nr1ps nuclear receptors.

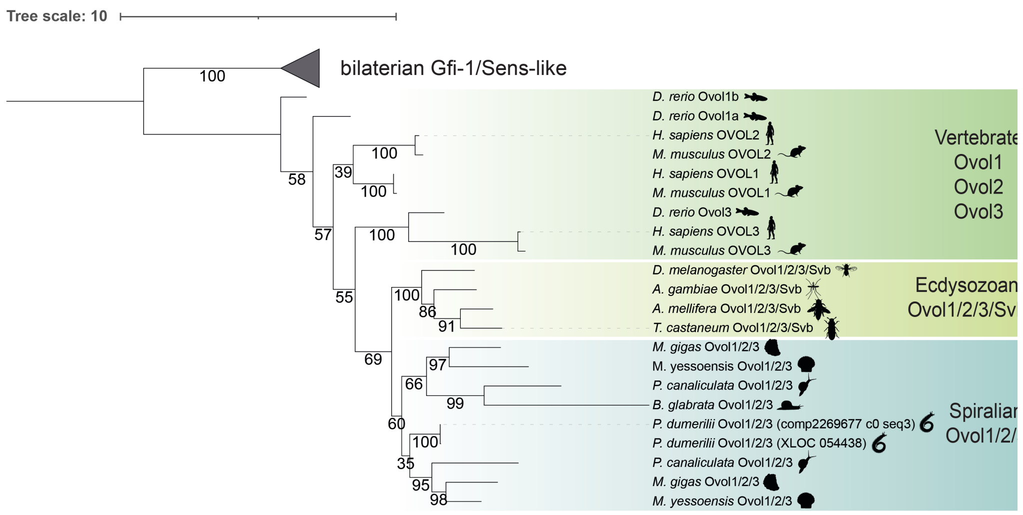


**figure S18:** Phylogenetic tree reconstruction of bilaterian Ovol1/2/3 zinc-finger transcriptional repressors.

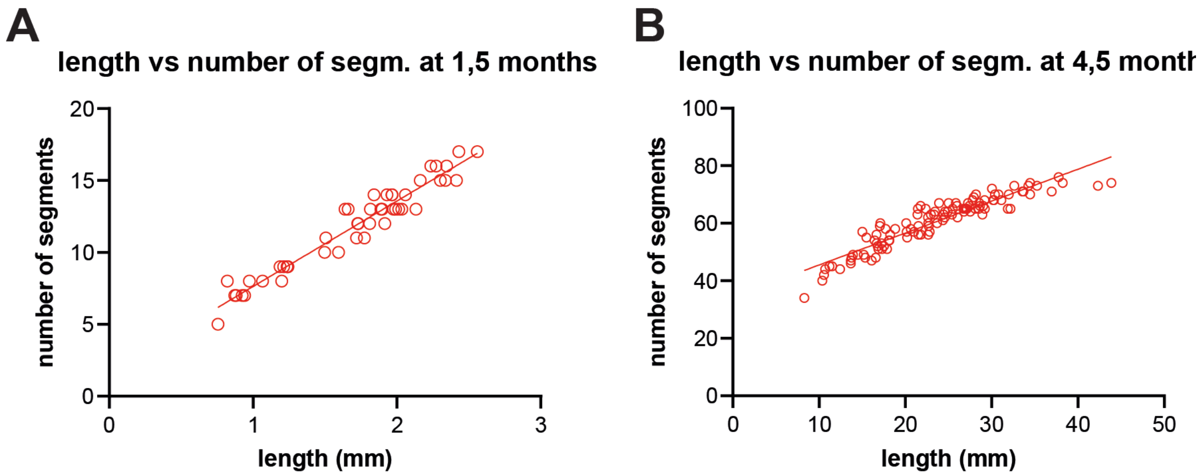


**figure S19:** Correlation between segments number and worm´s length in age-matched *l-cry-/-* mutants at 1,5 and 4,5 months. N ≥ 47 for each developmental stage. Simple linear regression, $R^2 = 0.9396$; $R^2 = 0.8617$, respectively.

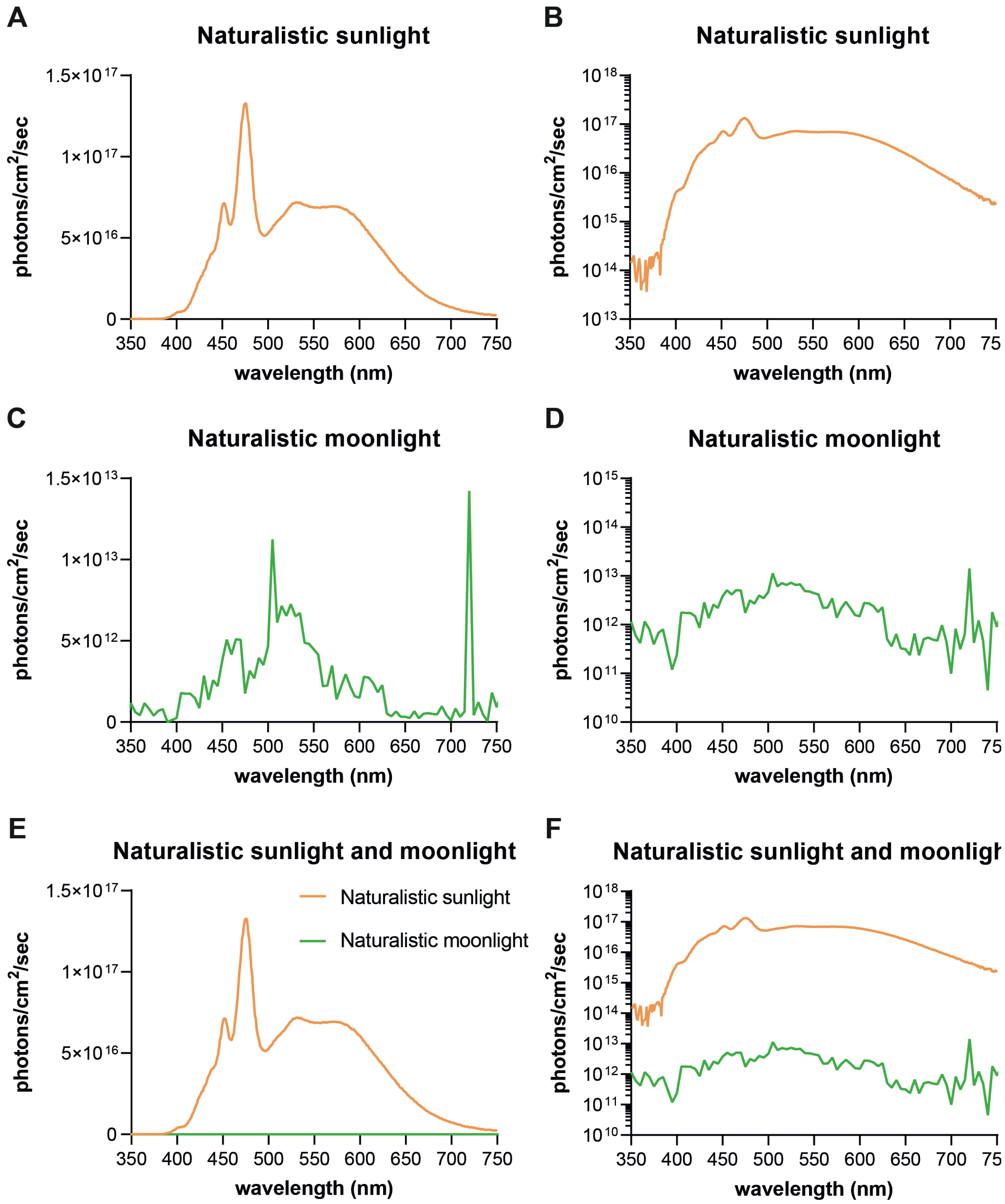


**figure S20:** Spectra of naturalistic sunlight and moonlight used in this study (experiment shown in Fig. 2 and fig. S2; LDM: both sunlight and moonlight; LL: only sunlight). **(A, B)** Spectra of naturalistic sunlight (orange), (A) linear, (B) logarithmic ($log_{10}$) plots. **(C, D)** Spectra of naturalistic moonlight (green), (C) linear, (D) logarithmic ($log_{10}$) plots. For moonlight, photons flux values were averaged every 5 nm range. **(E, F)** Spectra of naturalistic sunlight (orange) and moonlight (green), (E) linear, (F) logarithmic ($log_{10}$) plots. For moonlight, photons flux values were averaged every 5 nm range.

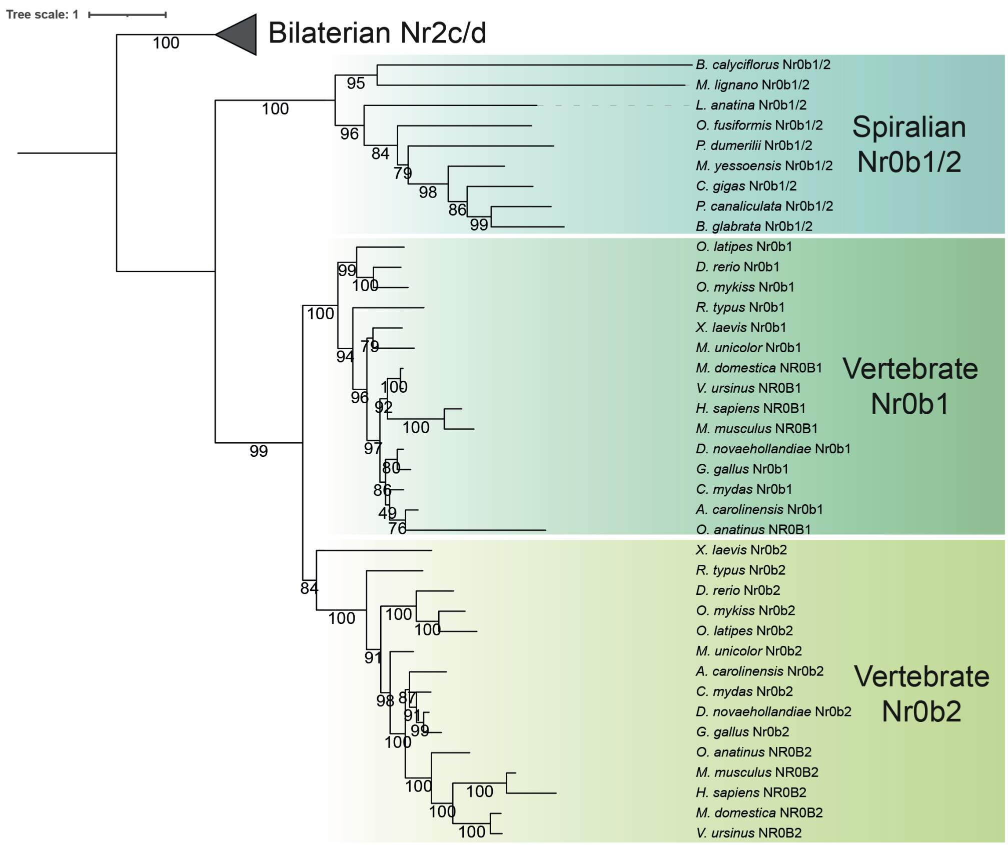


**figure S21:** Extended phylogenetic tree reconstruction of bilaterian Nr0b1/2 nuclear receptors including all sequences used for the alignment shown in fig. S11.

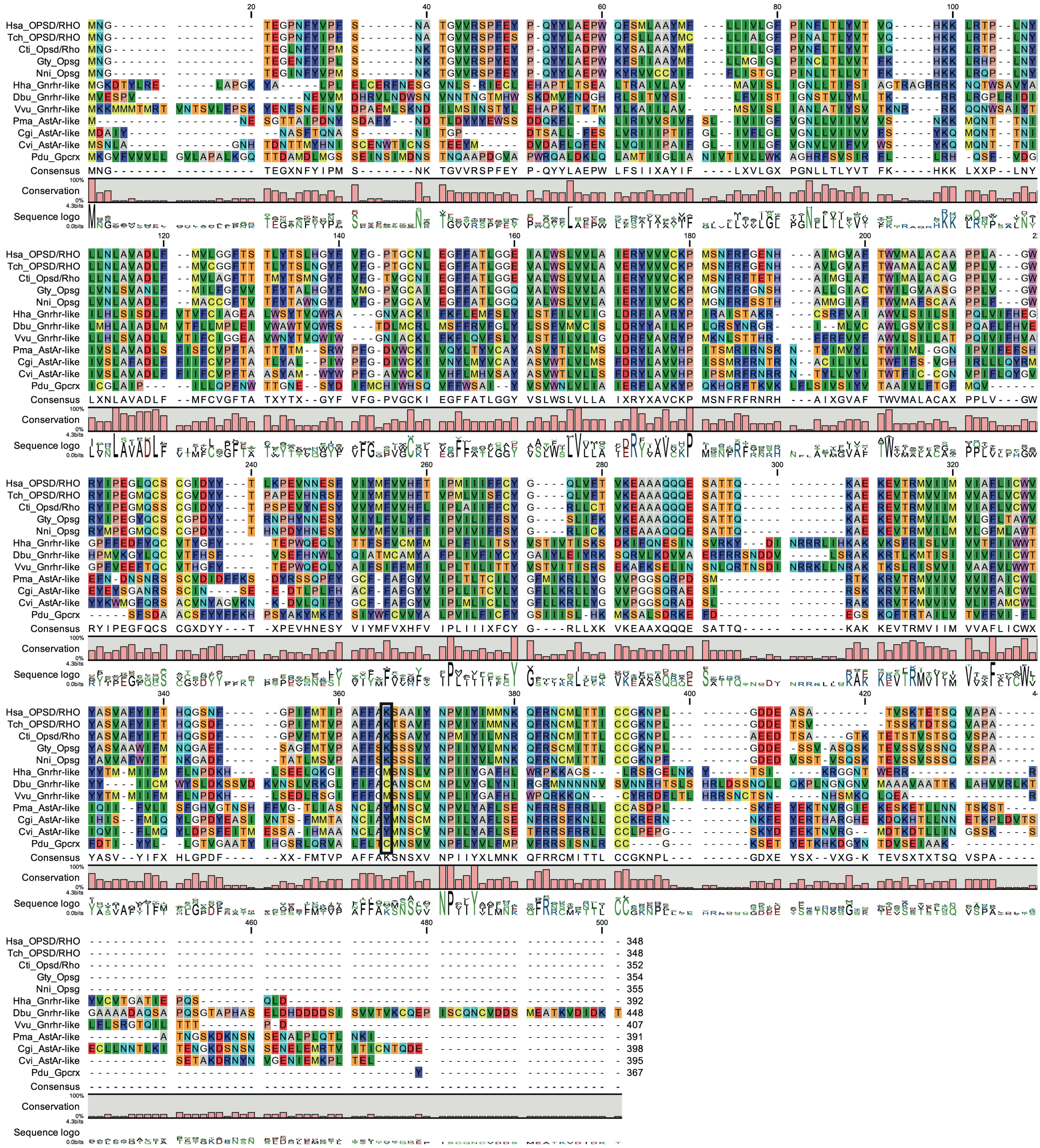

**figure S22:** Alignment of representative best BLAST hits for *Platynereis* Gpcrx using vertebrate, insect and mollusc databases. The black rectangle highlights the amino acid position at which the Lys necessary to form the Schiff base is present in light-sensitive GPCRs.

### Supplementary Table legends

**Table S1:** Differential expression analyses between *l-cry-/-* vs *l-cry+/+* immature and premature female worm heads. Related to Fig. 3A-E.

**Table S2:** Differential expression analyses between *l-cry-/-* vs *l-cry+/+* and *nr0b1/2+/-* vs *nr0b1/2+/+* immature worm heads. Related to Fig. 5D-F, H; fig. S4B, E, G.

**Table S3:** Differential expression analyses between *l-cry-/-* vs *l-cry+/+*, *nr0b1/2-/-* vs *nr0b1/2+/+*, *nr0b1/2+/-* vs *nr0b1/2+/+*, and *nr0b1/2-/-* vs *nr0b1/2+/-* immature worms. Related to Fig. 5A-C, G, H; fig. S4A, C, D, F.

**Table S4:** Summary tables of major candidate pathways and genes identified in the differential expression analyses between *l-cry-/-*, *nr0b1/2-/-*, and *nr0b1/2+/-* genotypes vs respective wt controls, and considered in this study. Related to Fig. 3A-E, Fig. 5A-I; fig. S4A-H.

**Table S5:** Best hits obtained blasting *gpcrx* transcript (initially annotated as *Green-sensitive opsin*, *OPSG*) against vertebrates, insects, and molluscs protein databases.

**Table S6:** Best hits obtained blasting all transcript sequences grouped as *gpcry* (all initially annotated as *OPSD/Rhodopsin*) against vertebrates, insects, and molluscs protein databases.

**Table S7:** Protein sequences (and correspondent annotations and accession numbers) used for all phylogenetic tree reconstructions and alignment in this study.

**Table S8:** Whole temperature and light recordings from HOBO devices placed to monitor environmental conditions under the different light regimes used to evaluate light-dependent effects on *Platynereis* lifespan and growth. Related to Fig. 2 and fig. S2.

**Table S9:** Statistical analyses of lifespan data presented in this work. Related to Fig. 1B; fig. S1A; Fig. 2B; fig. S2A; Fig. 4C; fig. S3A.
